# Flexible Circuit Mechanisms for Context-Dependent Song Sequencing

**DOI:** 10.1101/2021.11.01.466727

**Authors:** Frederic A. Roemschied, Diego A. Pacheco, Elise C. Ireland, Xinping Li, Max J. Aragon, Rich Pang, Mala Murthy

## Abstract

Many sequenced behaviors, including locomotion, reaching, and vocalization, are patterned differently in different contexts, enabling animals to adjust to their current environments. However, how contextual information shapes neural activity to flexibly alter the patterning of actions is not yet understood. Prior work indicates such flexibility could be achieved via parallel motor circuits, with differing sensitivities to sensory context [1, 2, 3]; instead we demonstrate here how a single neural pathway operates in two different regimes dependent on recent sensory history. We leverage the *Drosophila* song production system [4] to investigate the neural mechanisms that support male song sequence generation in two contexts: near versus far from the female. While previous studies identified several song production neurons[5, 6, 7, 8], how these neurons are organized to mediate song patterning was unknown. We find that male flies sing ‘simple’ trains of only one syllable or mode far from the female but complex song sequences consisting of alternations between modes when near to her. We characterize the male song circuit from the brain to the ventral nerve cord (VNC), and find that the VNC song pre-motor circuit is shaped by two key computations: mutual inhibition and rebound excitability [9] between nodes driving the two modes of song. Weak sensory input to a direct brain-to-VNC excitatory pathway (via pC2 brain and pIP10 descending neurons) drives simple song far from the female. Strong sensory input to the same pathway enables complex song production via simultaneous recruitment of P1a neuron-mediated disinhibition of the VNC song pre-motor circuit. Thus, proximity to the female effectively unlocks motor circuit dynamics in the correct sensory context. We construct a compact circuit model to demonstrate that these few computations are sufficient to replicate natural context-dependent song dynamics. These results have broad implications for neural population-level models of context-dependent behavior [10] and highlight that canonical circuit motifs [11, 12, 13] can be combined in novel ways to enable circuit flexibility required for dynamic communication.

## 1 Main

During courtship, *Drosophila* males chase and sing to females [14, 15, 16, 17]; male song is generated via wing vibration and composed of two main modes termed ‘pulse’ and ‘sine’. Male song patterns, timing, and intensity are known to be modulated by feedback cues stemming from the female [18, 19]. Here, we sought to investigate how song production neurons [5, 6, 7, 8, 20, 21, 22] are functionally organized to generate different song patterns in different sensory contexts. We utilize a combination of broad-range optogenetic activation in freely behaving animals, automated behavioral quantification, neural recordings and manipulations, and circuit modeling, and focus on four cell types in the brain and ventral nerve cord known to be sexually dimorphic and to contribute to singing. These include: P1a courtship arousal promoting neurons [23, 24, 25], pC2 neurons [8, 26], pIP10 descending neurons [5], and TN1 VNC neurons [6].

## 2 *Drosophila* males alter song sequence composition near versus far from the female

During courtship, *Drosophila melanogaster* males compose their songs into bouts interleaved with silences - each song bout consists of either simple trains of a single mode (‘pulse’ or ‘sine’ only) or complex trains of rapid alternations between song modes (Fig. 1a-b), and males continually switch between singing these different types of song throughout courtship (Fig. S1a). Prior work demonstrated that males produce pulse song, the louder mode, at larger distances to the female, and sine song once closer [18, 19, 20]. We collected a large data set of courtship interactions, combining both high-resolution video and audio recordings throughout a 36 mm behavioral chamber (or 12 fly body lengths; [18, 27]). When examining song bout composition, we find an abrupt change at approximately 4mm (or roughly 1.3 fly body lengths) distance: in close proximity of the female, males sing longer, complex bouts composed of alternations between pulse and sine elements, but beyond 4mm they sing shorter pulse-only bouts (Fig. 1c-e; Fig. S1b); these two contexts also correspond to differences in male forward velocity (mFV; Fig. S1e-f). We term these two contexts ‘near’ and ‘far’ throughout the study. Increasing bout complexity may be desirable to the female, as the majority of bouts immediately preceding copulation are complex (Fig. S1c-d).

**Figure 1:**
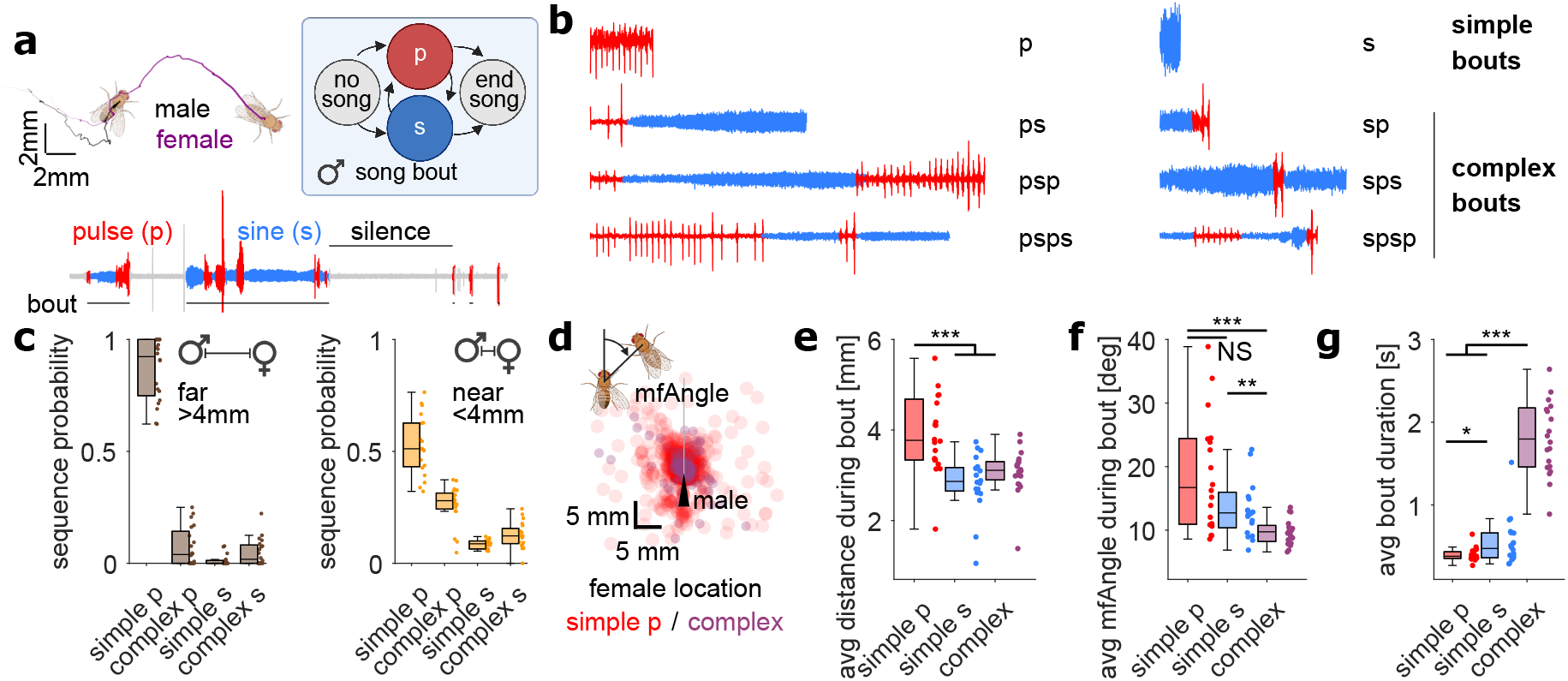
Context-Dependent Differences in Song Sequencing in *D*rosophila melanogaster. **a**, *Drosophila* male courtship song is structured into bouts comprising two main modes, ‘pulse’ and ‘sine’. **b**, Song bouts consist of either simple pulse (p) or sine (s) trains, or complex sequences that involve continuous alternations between modes. **c**, The distribution of song sequence types is different far versus near the female. Complex p = all complex bouts that start in pulse mode. Complex s = all complex bouts that start in sine mode. **d**, Location of the female (circles) during the production of male song (simple pulse (red) vs. complex bouts (purple)), in male-centric coordinates (male is depicted by the black triangle). Male-female angle (mfAngle) is the angle of the female thorax relative to the male’s body axis. Complex bouts are more likely to be produced when females are close and directly in front of the male. **e**, Average male-female distance during simple and complex song bouts. **f**, Average male-female angle during simple and complex song bouts. **g**, Average duration of simple and complex song bouts. **c-g**, n=20 wild type males (courting wild type females) - see Table S2 for genotypes **e-g**, Wilcoxon rank-sum test for equal medians; *P < 0.05, **P < 0.01, ***P < 0.001, 1; NS, not significant.

Intriguingly, song at all distances is biased to bouts with leading pulse song (‘p’ for pulse-only bout or ‘ps…’ for complex bout starting with pulse). Both far from and near the female, simple pulse bouts (‘p’) constituted the majority of all bouts (> 95% and around 55%, respectively), followed by complex ‘ps…’ bouts near the female (around 30%; Fig. 1c). Simple sine bouts (‘s’) constituted the minority of bouts at all distances. This apparent bias for bouts with leading pulse song suggests that the song pathway is organized to drive activity in pulse-generating neurons initially, in both contexts. The production of complex sequences might then arise via reciprocal interactions between pulse and sine producing neurons, but only in the near context. Finally, as the change in song complexity near the female is coupled with longer song bouts (Fig. 1e,g), inhibition to the song pathway (to suppress song when no female is present, or to keep song bouts short when far from a female) may be lifted when the male is near the female.

Taken together, these behavioral results favor a model in which context-dependent changes in song complexity arise through modulation (including a disinhibitory motif) of a single pathway. To test these hypotheses about song circuit organization, we used optogenetic activation of key song producing neurons coupled with quantitative analysis of song behavior.

## 3 A VNC rebound circuit enables generation of complex song bouts in the presence of a female

We expressed the red-shifted channelrhodopsin CsChrimson [28] in two types of song-producing neurons, either pIP10 brain-to-VNC (ventral nerve cord) descending neurons (1 neuron per hemisphere, Fig. 2a; [5]) or TN1 VNC neurons (a population of roughly 30 neurons in the wing neuropil of the VNC divided into 5 subtypes, TN1A-E, Fig. 2a; [6]), and analyzed song produced [17] following bilateral activation. Even though *Drosophila* males sing via unilateral wing vibration, both the extended wing and the closed wing receive the identical motor activity during song production [29]. We focused on these two cell types because previous work found that pIP10 and TN1A are primary drivers of pulse and sine song, respectively, suggesting that the production of complex bouts (containing a mix of pulse and sine) may be coordinated by neurons upstream that toggle between two pathways, one driving pulse song via pIP10 and the other driving sine song via TN1A [30]. By utilizing an optogenetic stimulation protocol that spanned multiple orders of magnitude in both irradiance and duty cycle (randomized full factorial design; Fig. 2a-b), we explored how varying activity in these two cell types, each critical for one mode of song, affected production of the other mode; critically we analyze behavior (here song) at sufficiently high resolution to relate neural activity to behavior. We hypothesized that if the pulse and sine pathways are interconnected, we might observe sine song following pulse song activation and pulse song following sine song activation. Such interactions may have been missed in prior work if they required activation outside the selected stimulation regime, or were under inhibition in the chosen experimental conditions (e.g., activation in isolated males).

**Figure 2:**
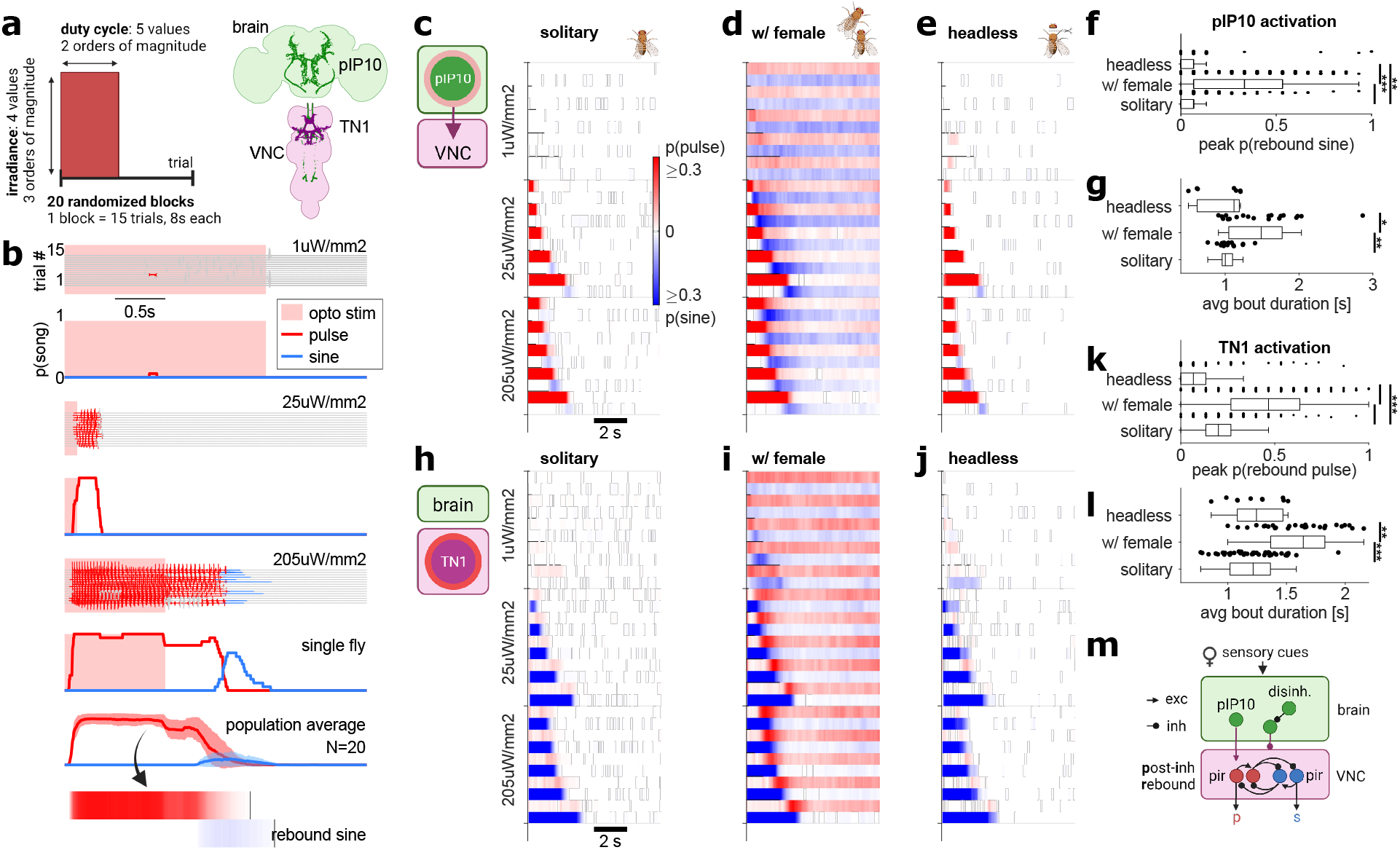
Reciprocal interactions between pulse- and sine-producing neurons in the presence of a female. **a**, Broad-range optogenetic stimulation paradigm involves systematic variation in optogenetic stimulus irradiance and duty cycle to map the relationship between neural activity in song neurons (pIP10 descending neurons and TN1 VNC neurons; schematic based on data from [6, 21]) and song production dynamics (see Methods for details). **b**, Song production on each trial and time-resolved song probabilities across trials following optogenetic activation of pIP10 neurons in a solitary male. Responses are shown for three out of 20 randomized stimulus blocks. At bottom, the population-averaged pulse and sine probability for the third example stimulus block (*n* = 20 recordings). The production of sine song immediately following pulse song production is indicated and termed ‘rebound sine’. **c-e**, Average song probabilities (see b) for all stimulus blocks (each pair of rows is a distinct optogenetic stimulus) and following optogenetic activation of pIP10 in solitary males (c), males paired with a wild-type female (d), and decapitated solitary males (e). **f**, Quantification of rebound sine probability following activation of pIP10 neurons (for the highest irradiance level) in solitary, female-paired, or headless males. The presence of the female promotes complex bout (pulse followed by rebound sine) generation following pIP10 activation. **g**, Average duration of song bouts generated via optogenetic activation of pIP10 neurons in solitary, female-paired, or headless males. The presence of the female promotes longer song bouts following pIP10 activation. **h-j**, Average song probabilities (see b) for all stimulus blocks (each pair of rows is a distinct opto stimulus) and following optogenetic activation of TN1 neurons in solitary males (h), males paired with a wild-type female (i), and decapitated solitary males (j). **k**, Quantification of rebound pulse probability following activation of TN1 neurons in (for the highest irradiance level) solitary, female-paired, or headless solitary males. The presence of the female promotes complex bout generation (sine followed by rebound pulse) following TN1 activation. **l** Average duration of song bouts generated via optogenetic activation of TN1 neurons in solitary, female-paired, or headless males. The presence of the female promotes longer song bouts following TN1 activation. **m**, Circuit model of song pathway based on broad-range optogenetic activation results: cues from the female ‘unlock’ complex bout generation via modulation of post-inhibitory rebound excitability in pulse and sine driving neurons of the VNC. **c-g**, n=20/20/10 males. For h-i, n=23/28/10 males. **f,g,k,l**, Wilcoxon rank-sum test for equal medians; *P < 0.05, **P < 0.01, ***P < 0.001, 1; NS, not significant.

Consistent with previous findings (cf. [5, 6, 21]), activation of either pIP10 or TN1 neurons in solitary males predominantly drove stimulus-locked pulse or sine song, respectively (Fig. 2c,h). However, we found that strong optogenetic stimuli drove pIP10 neurons to produce ‘rebound’ sine song following the offset of optogenetically-elicited pulse song (Fig. 2b-c; Fig. S2a). Strong stimulation of TN1 neurons drove reliable sine song with a small proportion of intermittent pulse song (Fig. 2h; Fig. S2b), as expected given that the TN1 genetic driver line (see Methods) labels some pulse song-producing VNC neurons [6].

The restriction of rebound sine following activation of pIP10 to high optogenetic activation levels indicates that the activity dynamics that generate complex bouts (pulse-sine alternations) are under inhibition in solitary males, possibly due to a lack of male arousal. In support of this idea, baseline inhibition masking latent excitatory pathways has already been identified in the *Drosophila* VNC, and is hypothesized to support context-dependent gating of proprioceptive information [31, 32]. Consistent with this hypothesis, optogenetic activation of either TN1 or pIP10 in males paired with females reliably drove long bouts of complex song across a broader range of stimulus parameters versus in solitary males (Fig.2d,f-g,i, k-l, Fig. S2c-d) and this complex song was driven predominantly when near the female (Fig. S2e-f), suggesting that female sensory cues functionally disinhibit the song pathway, unlocking the ability of pIP10 or TN1 neurons to produce complex song.

The observed dependence of rebound song on female sensory cues (not present when pairing activated males with a wild-type male, Fig. S2g-h) could arise via activation of P1a neurons known to mediate male arousal close to (and driven by tapping of) the female (Fig.2m;[33, 34]). Mutual inhibition between pulse and sine producing neurons (red and blue nodes in Fig.2m), combined with cell-intrinsic rebound excitability [9], could account for complex bout generation in the disinhibited circuit: activity of the pulse nodes would produce pulse song and inhibit sine production, whereas termination of activity in the pulse nodes would stop pulse song production and release inhibition of the sine nodes, leading to post-inhibitory rebound activity and production of rebound sine song. In this simplistic model, the pulse and sine nodes consist of two units: one that provides excitation (to drive motor output) and and another that provides inhibition (to suppress the other song mode). Inhibitory connections between the two inhibitory nodes, in addition to each excitatory node, would prevent premature bout termination - we provide further evidence for the existence of inhibitory connections between inhibitory neurons below.

If (part of) the proposed rebound circuit resides in the brain, pIP10 or TN1 neuronal activation would not be able to drive rebound dynamics (complex song) following decapitation. However, we found that activation of either pIP10 or TN1 neurons in decapitated male flies resulted in rebound sine or pulse song, respectively, comparable to that produced in intact males (Fig.2e-f,j-k, Fig. S2i-j), suggesting the rebound circuit exists in the VNC.

## 4 VNC song neurons display signatures of mutual inhibition and rebound excitability

We next examined neural activity in the VNC for signatures of the proposed rebound circuit (Fig. 2m). We optogenetically activated pIP10 neurons while simultaneously recording from neurons expressing the sex-specific transcription factor Doublesex (*D*sx) in the VNC via two-photon calcium imaging (Fig. 3a-b). VNC Dsx+ neurons include TN1 neurons [6] and dPR1 neurons [5], previously implicated in sine song and pulse song production, respectively, and all are excitatory [26]. While TN1A neurons drive sine song, other subtypes of Dsx+ TN1 neurons likely contribute to pulse song production [6]. We therefore expected to observe neural activity among the TN1 population both correlated with pIP10 activation (and therefore implicated in pulse song production) or anti-correlated with activation (and therefore implicated in rebound sine song production) - importantly these subsets should be distinct from each other across repeated optogenetic stimulation.

**Figure 3:**
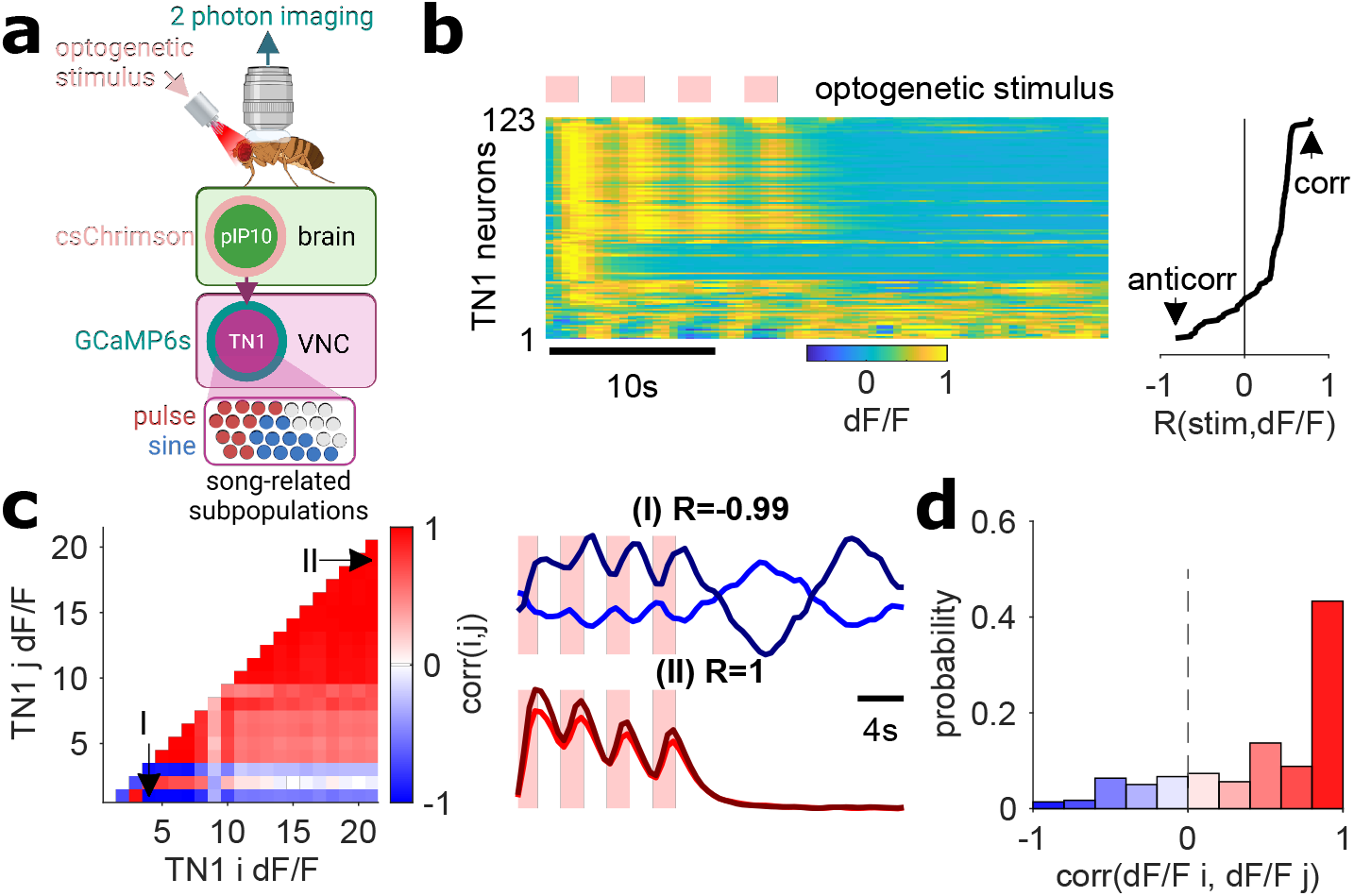
Calcium imaging from Dsx+ TN1 neurons of the ventral nerve cord. **a**, Two-photon calcium imaging from VNC Dsx+ TN1 neurons combined with optogenetic activation of pIP10 descending neurons (see Methods for details; see Table S2 for genotypes). **b**, TN1 neurons show diverse calcium response dynamics following pIP10 activation (123 TN1 neurons across 3 flies). Each row shows the normalized calcium response (dF/F) of a single soma, averaged over 7 trials (and each trial containing four stimulus presentations), and responses are sorted by their correlation with the optogenetic stimulus. The optogenetic stimulus pattern was chosen to produce pulse song followed by rebound sine (see Fig. 2c) in solitary males. **c**, Pairwise correlation of trial-averaged activity between any two TN1 neurons from one male, following pIP10 activation. Examples of strong anti-correlation and correlation are shown at right (I/II; P<1e-50). **d**, Distribution of calcium response correlation coefficients across TN1 recordings in *n* = 3 males. Colors as in (c).

We used an optogenetic protocol that combined long stimuli driving temporally separated pulse and rebound sine when activating pIP10 in solitary, freely behaving males (Fig. 2c, both intact and headless males), with short inter-stimulus-intervals to maximize the magnitude of evoked calcium responses, making it possible to observe neural activity associated with both pulse and sine song production. We observed a broad range of temporal response patterns within the TN1 population following pIP10 activation. The activity of nearly half of the recorded TN1 neurons was positively correlated to the pIP10 stimulus, while a smaller fraction showed activity anti-correlated to the stimulus, activity only to the first stimulus, or intermediate to no correlation (Fig. 3b). Imaging from pIP10 axons and Dsx+ dPR1 neurons showed only tight correlation to the optogenetic stimulus (data not shown).

By examining pairwise activity correlations in TN1 neurons, we found that a subset of pairs exhibited nearly perfect anti-correlation in their responses to pIP10 activation, with alternating peaks in activity persisting beyond the stimulation (Fig. 3c-d, Fig. S3a-b), as expected from neurons that are both (functionally) mutually inhibitory and rebound excitable [9]; we found these anti-correlated pairs on both sides of the VNC (as expected, to drive pulse-sine rebound activity in both wings). We had expected to detect only a small number of sine-producing neurons, given the fraction of stimulus presentations with rebound sine rarely exceeded 30% in solitary or headless males (Fig. 2c,e,f). We hypothesize that this anti-correlated subset consists of the TN1A sine-producing pre-motor neurons [6] - since TN1 neurons are known to be excitatory (being part of the cholinergic neural cluster of hemilineage 12A [26, 35]), and we hypothesize the existence of intermediary inhibitory neurons responsible for coupling in the rebound circuit (Fig. 2m). Due to the near-perfect anti-correlation observed for some neuron pairs, we assume that these intermediary neurons are mutually inhibitory, as indicated in (Fig. 2m), in addition to providing inhibition to song-generating neurons, thus implementing an ‘adaptive switch’ circuit motif with a known role in stimulus categorization [36, 37].

In conclusion, VNC TN1 neurons exhibit hallmarks of a rebound circuit that could underlie complex song dynamics. We next ask how neural activity in the brain drives or modulates this rebound VNC circuit to produce the observed sensory gating and context-dependence of complex song.

## 5 Acute sensory cues from the female both drive pulse song production and enable complex song bouts

Having uncovered a rebound circuit in the VNC, we next asked what brain mechanisms drive this circuit. We again used our broad-range optogenetic activation protocol (see Fig. 2a) and explored the role of P1a [7, 23] and pC2 [8, 26, 38] brain neurons in song production - these cell types were previously implicated in male song production, but their role relative to song sequencing was unknown.

### pC2 neurons simultaneously drive pIP10 descending neurons and P1a neuron-mediated disinhibition

P1a neurons are primarily activated by taste cues collected when males tap females during courtship [34]; these neurons in turn can drive a persistent arousal state via downstream recurrent circuitry [39]. Because P1a neurons have been suggested to be upstream of pIP10 neurons [5], we hypothesized that activating P1a neurons in solitary males would mimic our results with pIP10 activation in the presence of a female (Fig. 2d). In contrast, we found that activation of P1a neurons (again using our broad optogenetic stimulation paradigm) in solitary males produced persistent and variable song but lacked the stereotyped stimulus-triggered complex song sequences observed for activation of pIP10 neurons in the presence of a female (Fig. 4a, Fig. S4a), suggesting that P1a activity alone is insufficient for temporally precise control of song bout complexity.

**Figure 4:**
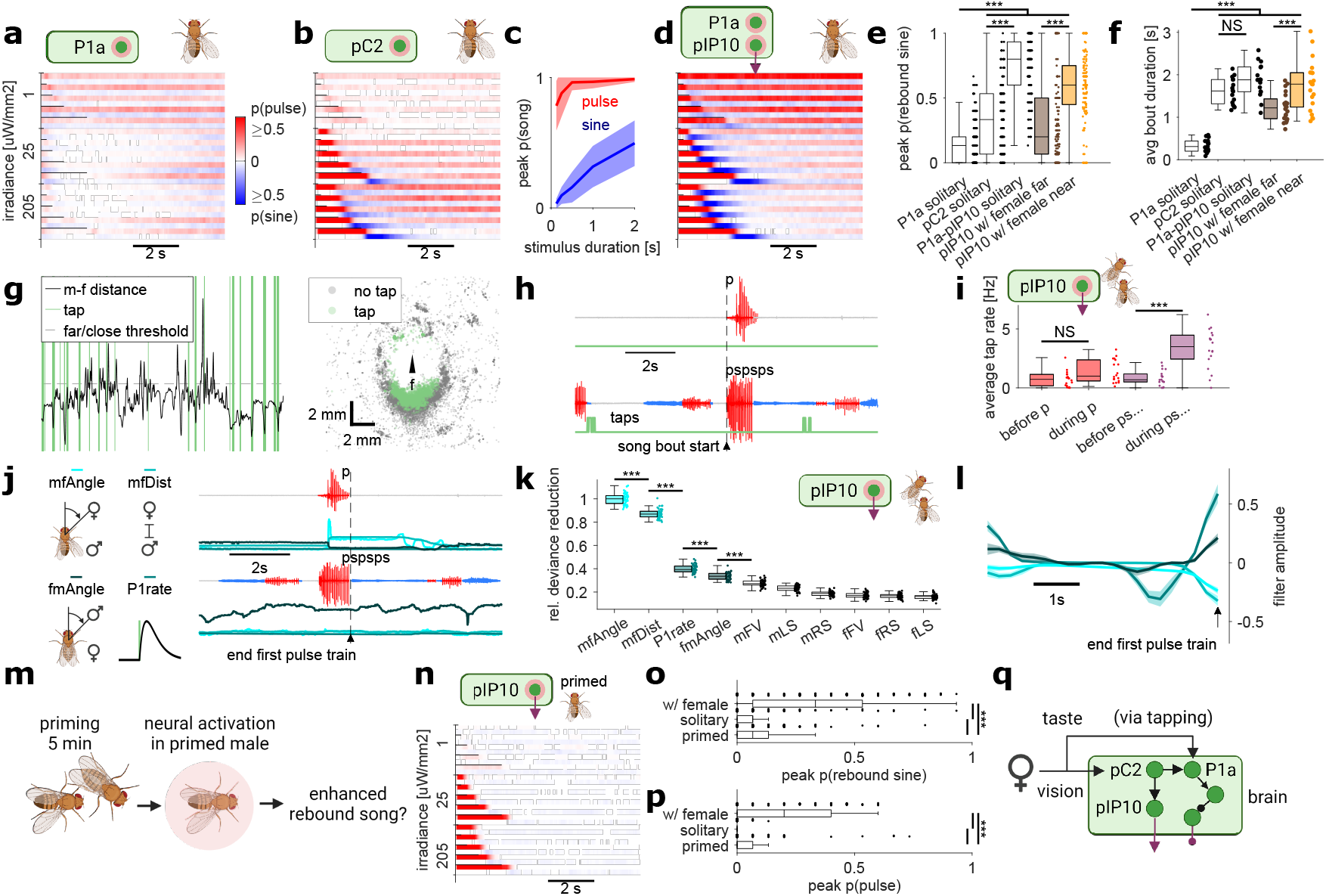
Acute female sensory cues promote complex song bout generation. **a,b,d**, Quantification of pulse and sine song probabilities following optogenetic activation of P1a (a), pC2 (b), or both pIP10 and P1a neurons (d), in solitary male flies (*n* = 17/16/16). See Table S2 for genotypes. **c**, Peak pulse and sine song probability as a function of stimulus duration for intermediate-irradiance activation of pC2 neurons (25uW/mm2). **e**, Comparison of peak rebound sine probability for intermediate- and high-level optogenetic activation (25 and 205 *μ*W/mm2) of different song neurons (pC2, pIP10 or P1a) in solitary males (data shown in (a,b,d)) as well as activation of pIP10 in males paired with a wild-type female (data shown in Fig. 2d). **f**, Comparison of average bout duration for groups and activation levels shown in (e). **g**, (left) Automated tap detection using a convolutional neural network (see Methods). Taps (green) detected in an example recording overlaid on male-female distance (black) and the 4mm threshold used for defining the far vs. near context (grey). (right) Male locations during tap (green) and no tap (grey) events, in female-centric coordinates, corresponding to the recording shown at the left. **h**, Examples of simple (top) and complex (bottom) pulse bouts along with detected taps (green raster). Tap rate was quantified as the number of taps within a song bout, divided by the duration of the bout (to compare with time before a bout, we used the number of taps within an equally-sized window preceding the bout, divided by bout duration). **i**, Average tap rate before and during simple and complex bouts with leading pulse song, for bouts driven by optogenetic activation of pIP10 in males paired with a wild-type female (Fig. 2d, *n* = 18). **j**, To fit generalized linear models (GLMs) predicting bout type (simple pulse vs. complex pulse), we used movement or interaction features or P1 rate (see Methods; shades of cyan) over the 5s preceding the end of the first pulse train within simple (top) or complex (bottom) pulse bouts (see Methods). **k**, Relative deviance reduction (from the GLM) for features predicting simple vs. complex bout production following pIP10 activation with a female. Input features are ranked by their predictive power (*n* = 51 fits of the model on random subsets of data from *n* = 18 recordings; see Methods). **l**, GLM filters are shown for the four most predictive features in (k). **m**, To test for effects of persistent male arousal on optogenetically driven song, males were primed (allowed to court a virgin wild-type female) for 5 minutes preceding optogenetic activation. **n** Song probabilities for optogenetic activation of pIP10 neurons in solitary males that were primed. *n* = 19. **o**, Comparison of peak rebound sine probability for optogenetic activation of pIP10 following priming at intermediate and strong irradiance (25 and 205uW/mm2). **p**, Comparison of peak pulse probability for optogenetic activation at lowest irradiance (1uW/mm2) of pIP10 in primed, solitary, or female-paired males. Groups identical to those in (o). **q**, Updated model of the male brain circuitry involved in context-dependent song sequencing. **j-l**, mfAngle = male-female angle, mfDist = male-female distance, fmAngle = female-male angle **e,f,i,k,o,p**, Wilcoxon rank-sum test for equal medians; *P < 0.05, **P < 0.01, ***P < 0.001, 1; NS, not significant.

pC2 neurons in males [40] are known to be responsive to both visual and auditory cues [8, 38] - additionally, in the female hemibrain [41], pC2 neurons receive direct inputs from both visual (lobula columnar cells) and auditory projection neurons. We found that activation of pC2 neurons in solitary males robustly drove pulse song followed by rebound sine, similar to pIP10 activation in the presence of a female (Fig. 4b, Fig. S4b, Fig. S5a-b). Interestingly, we also observed persistent and variable song in the period outside of optogenetic activation of pC2 neurons in solitary males. Notably, for intermediate optogenetic irradiances that reliably drove pulse song, the amount of rebound sine following pC2 activation showed a near-linear dependence on the duration of the optogenetic stimulus (Fig. 4c), suggesting that distinct levels of pC2 neural activity control the transition from simple to complex bout generation. Pulse song is also composed of two main pulse types (termed Pfast and Pslow; [20]). Intriguingly, the duration of pC2 activity also determined the selection of pulse type, since brief activity mainly drove Pfast (as did activation of pIP10 in solitary males), whereas more sustained activity increased the relative amount of Pslow (as did activation of pIP10 when near a female; S4c). These results support the conclusion that pC2 neurons serve as the main determinant of song composition.

Taken together, these results suggest that pC2 neurons directly drive pulse song production via pIP10, but simultaneously drive P1a neurons to generate persistent song and unlock or disinhibit the rebound circuit in the VNC to enable complex song bouts (Fig. 4q). In line with this model, simultaneous activation of pIP10 and P1a neurons in solitary males produced highly reliable and long complex bouts, well beyond the levels observed for activation of the individual neuron types, including pIP10 activation in males near a female (Fig. 4d-f, Fig. S4d).

These results support the conclusion that P1a mediates disinhibition (via as of yet unknown circuitry) rather than excitation of the VNC circuit. While functionally similar, the former is computationally favorable, as disinhibitory gating preserves the dynamic range for processing of sensory information in target neurons, and it reduces spurious responses [42]. If P1a neurons instead mediated excitation of the VNC rebound circuit, we would expect to observe optogenetically-driven song bouts during activation of P1a in solitary males (which we do not). However, if P1a neurons disinhibit the rebound circuit, then a separate source of excitation is needed to drive song sequences. We have now identified that source as the pC2 neurons that mediate parallel drive to both pIP10 and P1a neurons (Fig. 4q).

### Identification of the sensory cues that distinguish simple from complex song bouts

During courtship, males respond to both visual and chemosensory cues from the female - changes in female locomotor speed and position drive changes in song production [18, 19] while taste cues collected via tapping of the female abdomen drive changes in arousal state via activation of P1a neurons and downstream recurrent circuitry [34, 39]. We investigated the contribution of both types of sensory stimuli via simultaneous tracking of male and female posture and locomotion (via SLEAP; [27]), combined with song analysis [17] during optogenetic experiments. We also built a deep learning-based detector for male tapping of the female abdomen (see Methods; Fig. 4g; Fig. S4e) and found that male tap rate is higher during complex song bouts versus either before these bouts or during or before simple bouts (Fig. 4h-i), suggesting that acute (tap-triggered) activation of P1a during an ongoing bout, rather than P1a-mediated arousal on longer timescales, promotes complex bout generation. This was the case both for optogenetically driven song and wild-type song (Fig. 4i; Fig. S4f), implying that P1a modulates behavior not only on the long timescale of persistent arousal [39], but also on the shorter timescale of sensory processing.

Yet, the fact that pC2 activation can drive complex song in solitary males (Fig. 4c) implies that sensory modalities other than tapping may also modulate bout complexity. To investigate this, we tracked visual cues provided by the female (e.g., her speed and relative position to the male) and then used both kinds of stimuli (female visual features and taste cues collected via tapping) to predict the production of simple versus complex song bouts, using generalized linear modeling (GLM; [18]; Fig. 4j-l; see Methods). Specifically, for bouts driven by optogenetic activation of pIP10 (that is, those bouts starting during optogenetic activation in males courting a wild-type female), we predicted whether a bout starting in pulse mode either terminated (and hence was a simple pulse bout) or became complex bout (by switching to sine mode). The GLM stimulus history used for predictions covered the five seconds leading up to the termination of the first pulse train of a bout, and hence was blind to bout type (simple or complex).

We found that reductions in the angle of the female’s body relative to the male’s body axis (mfAngle, cf. Fig. 1d), in addition to male-female distance (mfDist), within the one second leading up to the end of the first pulse train in a song bout, were the most predictive of whether the pulse bout ended (that is, remained simple) or continued to become a complex pulse-sine alternating bout (Fig. 4j-l). Compared to mfAngle, the relative predictive power (relative deviance reduction, see Methods) of an estimate of P1a activity derived from the tap detection data (see Methods) was roughly 40%, suggesting that tapping and the resulting activity of P1a neurons alone do not fully predict bout complexity. Nonetheless, changes in the estimated P1a activity from below-average to above-average within the two seconds prior to the end of the initial pulse train were predictive of a complex bout, in line with our previous analysis of tap rate (Fig. 4i). These results imply that combined sensory modalities contribute to song bout complexity (and these results also hold for wild-type song, Fig. S4g): primarily visual activity relayed through pC2 and tap rate relayed through P1a both contribute to driving complex song sequences (Fig. 4q). We test whether this sensory information is sufficient to generate naturalistic bout statistics below in our computational circuit model.

Notably, these sensory effects on song patterning act on a timescale of milliseconds to seconds that is much faster than the known role of P1a in predominantly driving minutes-long, persistent, changes in behavior [23, 39]. To test whether P1a-mediated persistent arousal also promoted complex song, we induced male arousal by allowing the male to court a virgin female during a five-minute ‘priming’ phase immediately preceding optogenetic activation (Fig. 4m). However, priming only weakly enhanced the complexity of optogenetically driven song in comparison to solitary males not subject to priming (Fig. 4n-o; Fig. S4h-j). However, we did observe that simple pulse song was induced in primed males even for the weakest levels of activation, in contrast to males not subject to priming, suggesting that male arousal modulates the excitability of pIP10 neurons at the timescale of minutes but without promoting complex song (Fig. 4p). These results corroborate that P1a neurons have a modulatory effect on behavior on a broader range of timescales than previously assumed.

Together, these results support a model (Fig. 4q) in which different sensory cues (vision or taste) contribute to the choice of simple vs. complex bouts: simple pulse bouts are driven by weak sensory input to pC2 (most likely visual since the most predictive features in the GLM were male-female angle and male-female distance) that propagates to the VNC via pIP10, whereas complex bouts can be driven either by strong input to pC2 which both activates pIP10 and P1a, or pIP10 activity combined with taste-driven P1a activity (Fig 4q). Having two mechanisms to drive P1a and downstream circuitry could facilitate continuous complex song production when a male transitions from following a moving female [25] to positioning himself behind a stationary female prior to copulation.

## 6 A circuit model with few canonical computations reproduces context-dependent song statistics in response to naturalistic input

Our behavioral and neural imaging results suggest that naturalistic song statistics arise from the specific functional architecture of the male song circuit. First, we find evidence for a core rebound circuit in the VNC with mutual inhibition between pulse and sine producing neurons and rebound dynamics in the inhibited nodes (Fig. 2m). Second, we find evidence for a direct pulse pathway from brain to VNC that integrates sensory signals from the female (Fig. 4q). Third, we find evidence for a disinhibitory brain pathway onto both nodes of the core circuit, which is driven by sensory input of different modalities (taste and vision via P1a and pC2, respectively). To test whether these few computational features are sufficient to explain naturalistic song statistics, we implemented them in a spiking neural circuit model that received naturalistic sensory input (dynamic changes in male-female distance) and of which the output were the spikes of the ‘pulse’ and ‘sine’ nodes of a core rebound circuit (see Methods; Fig. 5a).

**Figure 5:**
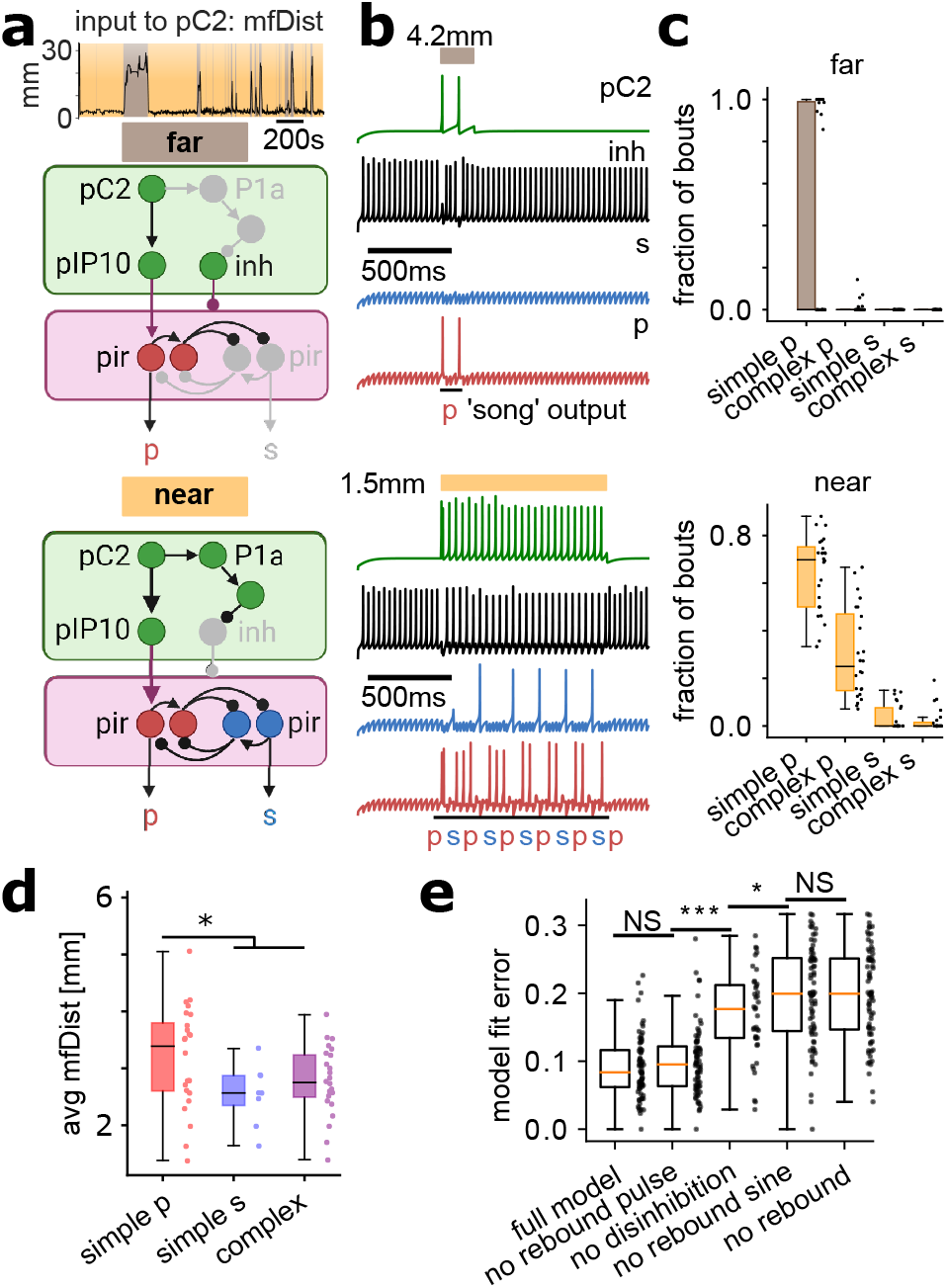
Neural circuit model of context-dependent song patterning. **a**, Neural circuit model of male song patterning far from and near a female. The only input to the model is fluctuating male-female distance (mfDist) taken from natural behavior recordings (top). This information enters the circuit at the level of the pC2 node (see Methods), which drives the pulse pathway. For strong input (when near the female), this input in parallel disinhibits the VNC rebound circuit to enable complex song production (alternating activity of the pulse and sine output nodes, p and s). Gray shading indicates parts of the circuit that become inactive at far or near conditions. **b**, Simulation of a spiking neuronal network comprising four nodes (pC2, inh, p, s; see Methods) that represent the key computations of the circuit in (a): disinhibition, rebound excitability, and mutual inhibition. Model parameters were fit to experimental data (from wild type courtship) using the genetic algorithm (GA) optimization. The resulting model was simulated using brief and weak (top, corresponding to mfDist=4.2mm) or long and strong (bottom, corresponding to mfDist=1.5mm) stimulation of the pC2 node, and produces (in this case) either simple (‘p’) and complex (‘psp…’) ‘song’ outputs. **c**, ‘Song’ statistics for *n* = 24 GA fits of a reduced version of the circuit in (a) to experimental song data (400 seconds each) at far or close distance (see Methods). The model, with only four nodes and three key computations, reproduces context-dependent bout statistics observed in courting wild-type flies (compare with Fig. 1c). **d**, Average male-female distance during simulated simple pulse, simple sine, or complex bouts (models same as in (c)) matches the relationship between distance and song types observed in courting wild-type flies (compare with Fig. 1e). **e**, Fit error (GA objective function) for the full model vs. models with individual computational features knocked-out (rebound excitability in the p, s, or both nodes, or disinhibition; see Methods). *n* = 72 (*n* = 39 for disinhibition knock out) 200-second long segments of song were randomly chosen from all wild-type recordings and the full and computationally reduced song circuit models were fit to each of these segments. **d,e**, Wilcoxon rank-sum test for equal medians; *P < 0.05, **P < 0.01, ***P < 0.001, 1; NS, not significant.

More specifically, the circuit model comprised four nodes (termed ‘pC2’,‘inh’ for inhibitory, ‘p’ for pulse, and ‘s’ for sine) for which the membrane dynamics were described by three ordinary differential equations (see Methods; [43]). Model parameters were chosen to facilitate tonic spiking in the pC2 and inh nodes and rebound spiking in the pulse and sine nodes (see Table S3). Sensory input to the pC2 node was modeled as naturalistic male-female distance (mfDist), which should account for both visual and taste inputs since males only tap when they are close to females. This input was then transformed into input current by a nonlinearity such that large/small mfDist corresponded to small/large input current to pC2 (see Methods and Table S3 for details). In the model, the inh node provided tonic inhibition onto the pulse and sine node, mimicking the male’s default, unaroused, state. pC2 provided excitatory input to the pulse node and functional inhibition to the inh node, mimicking the direct pulse pathway from pC2 via pIP10 to the VNC, and the proposed disinhibitory pathway from pC2 via P1a (activated for short mfDist and strong input to pC2; Fig. 5a), respectively.

The model had four free parameters: the gain of the sensory input current, the strength of the tonic input current to the inh node, and a global weight each for the inhibitory and excitatory connections. Under the simplifying assumption that spikes of the pulse and sine node correspond to pulse and sine song output, this allowed us to fit the model to experimental song statistics, using genetic algorithm optimization (see Methods). Surprisingly, our simple functional circuit model, comprising only four nodes that capture the essential computational features suggested by our data, was sufficient to recapitulate naturalistic song bout statistics both far from and near the female (Fig. 5b-d, cf. Fig. 1c,e).

Removing individual computations in the model (knocking out disinhibition, rebound excitability of the pulse, sine, or both nodes; see Methods) resulted in overall worse fits to the data compared to the full model (Fig. 5e), especially so when removing disinhibition or rebound excitability of the sine node. Notably, fit performance for a model lacking rebound pulse but capable of rebound sine was similar to that of the full model, highlighting the relative importance of rebound excitability of the sine node (compared to the pulse node) as a computational feature of the song circuit. This is consistent with our conclusion that the pulse production pathway is driven directly via sensory input to pC2 and subsequently to pIP10 (Fig. 4q), but that the sine node is not driven directly. An exclusively indirect drive of the sine node explains the small amount of simple sine song observed both in experiments and simulations (Figs. 1c and 5c), as disinhibition-mediated rebound activity can occasionally drive the sine neuron first (Fig. S6a), depending on the internal (membrane voltage) states of the sine and pulse neurons.

In our model, bout termination mainly relies on an increase in male-female distance (Fig. S6a-b) - this is consistent with analysis of natural courtship data that reveals that changes in female lateral speed (the female abruptly moving away from the male) are predictive of song bout termination ([18]). In addition, in the biological circuit other factors such as spike-frequency adaptation (present but not explicitly modeled here; cf. voltage trace of the ‘p’ node, Fig. 5b bottom) could play a role. In line with this hypothesis, we performed *in vivo* patch-clamp recordings of pIP10 and found clear signs of spike-frequency adaptation (Fig. S8a-c).

### Rebound circuitry in the VNC allows for complex bout control using a single command-like neuron

Our results suggest that the neural control of song generation is based on mutual inhibition between, and rebound excitability in, pulse and sine driving neurons in the VNC. While descending neuron pIP10 provides input to the pulse node, no equivalent descending neurons for sine song and hence no sine pathway is known, and our data and simulations suggest that such a descending sine pathway is not required to explain wild-type song bout statistics. One possible advantage of the proposed song circuit design based on a single descending (pulse) input to one node of a core rebound circuit is simplicity of control, since in principle the proposed circuit architecture allows for switching between simple pulse song and, theoretically, arbitrarily complex pulse-sine sequences, by simply adjusting the level and timing of pIP10 activity. Specifically, if pIP10 is activated at a level that drives rebound sine, and subsequently activated during every (rebound) sine train, this should generate an arbitrarily long sequence. To test this hypothesis, we used closed-loop optogenetic activation of pIP10 in a male courting a wild-type female, triggered on the real-time detection of sine song (see Methods and Fig. S7a-b). Indeed, such closed-loop activation significantly increased bout complexity and duration (Fig. S7c), confirming our hypothesis, and showing that in *Drosophila*, patterned activity of a single descending neuron (acting on a disinhibited VNC circuit due to female presence) suffices to generate highly complex song outputs.

### Descending activity from pIP10 is relayed via both electrical and chemical synapses, with distinct roles in song generation

Another direct prediction of the proposed circuit model is that blocking descending inputs to the core pulse node should strongly reduce the amount of bouts with leading pulse song. To test this prediction, we re-examined published data [20] with expression of tetanus toxin light chain (TNT,[44]) or inward-rectifying potassium channels (Kir2.1, [45]) in male pIP10 neurons to block chemical synapses originating in pIP10 or to tonically hyperpolarize pIP10, respectively. Since both TNT and Kir2.1 prevent chemical synaptic transmission, as expected, the amount of simple pulse-only bouts during courtship of a wild-type female was significantly reduced compared to genetic controls (Fig. S7d-e), suggesting that chemical synapses from pIP10 onto pulse-driving neurons downstream coordinate the execution of simple pulse song. However, males expressing TNT in pIP10 (but not Kir2.1) produced *more* simple and complex sine-leading bouts compared to controls (Fig. S7d-e), suggesting a role for electrical synapses (which remain intact in TNT flies) in mediating sine song generation. How could this occur? We posit that gap junction synapses between pIP10 and the inhibitory interneurons of the pulse-rebound circuit drive inhibition of sine pre-motor neurons (e.g., TN1A) when pIP10 is active, but when pIP10 activity ceases, inhibition of sine production is released, giving rise to rebound sine and inhibition of the pulse pre-motor neurons (Fig. S7f). This matches with our observation of near perfect anti-correlation between subsets of TN1 neurons (Fig. 3c-d; Fig. S3a-b). Thus, the re-evaluation of pIP10 inactivation data in light of our song circuit model suggests that song dynamics are shaped by an interplay of electrical and chemical synapses from pIP10 descending neurons onto the VNC rebound circuit.

## 7 Discussion

The ability to alter the sequencing of actions to match the current environmental context is observed across animals and behaviors, including for social interactions [46, 47, 48, 49, 50]. Here, we provide insights into the underlying circuit mechanisms by focusing on song production in two contexts in *Drosophila melanogaster*, near versus far from a female. Using quantitative behavior, modeling, broad-range optogenetics, circuit manipulations, and neural recordings, we find that simple song (of primarily the pulse mode) is driven by low-level/brief activation of pC2 brain neurons, which drive a pair of pIP10 brain-to-VNC descending neurons. To generate complex bouts, stronger, longer-duration pC2 neuron activity simultaneously drives pIP10 and recruits P1a neurons to functionally disinhibit core circuitry in the ventral nerve cord, allowing pIP10 descending signals to produce rapid alternations of pulse and sine song (complex bouts). Hence, the context-dependent execution of two different song modes (pulse and sine) does not require dedicated pathways for each mode, as had been previously postulated [30]. Rather, here the sensory context, encoded ultimately by acute P1a neural activity, determines which song repertoire is accessible to descending commands, effectively implementing context-dependence via two operational modes of a single circuit [51].

Context-dependence of acoustic communication is known in other species, including songbirds [52] and primates [53] - the circuit mechanisms we have uncovered here may therefore serve as a useful template in investigating those systems at the cellular level. Intriguingly, the presence of the female has opposing effects on song variability in flies and birds, species in which females prefer either variable [54] or stereotyped [55] song, respectively. In flies, we show that female proximity relieves the core song circuit from inhibition to promote song variability (rapid pulse-sine alternations of varying length), whereas in birds, female presence suppresses song variability via direct inhibition of basal ganglia neurons [56].

How do our results relate to prior work on song production and patterning in *Drosophila*? First, prior work suggested that pIP10 neurons drive only the pulse mode of song [5, 21] - however, those studies did not explore the broad range of optogenetic activation parameters used here, highlighting the value of varying neural activity levels during behavior to uncover circuit dynamics. Second, while our computational model of the song circuit can recapitulate song dynamics using only male-female distance as contextual information, prior work demonstrated that the male’s own locomotor speed is also highly predictive of song patterning: increases in male speed predict both pulse-leading bouts and transitions back to pulse song within a bout [18, 19]. This is consistent with our observation that males move faster in the ‘far’ context that consists primarily of pulse-only simple bouts (Fig. S1e-f). While we do not yet know where self motion information enters the song pathway, our model predicts that it should be integrated at the level of pIP10, pushing the song pathway towards pulse song production, without engaging the disinhibition arm of the pathway (via P1a neurons) that would lead to sine song production.

Third, prior work uncovered that there are two distinct types of pulse song termed Pfast and Pslow, and that the choice of pulse type depends on distance to the female [20]: males produce Pfast (the louder mode of song) at larger distances, switching to Pslow (the softer pulse type) when close. Our data indicates that the relative amount of Pfast and Pslow is ultimately controlled by the activity of brain pC2 neurons (S4c). Weak input to pC2 (when far from the female) drives weak activity in pIP10 and production of Pfast, while sustained activity of pC2 when near a female both functionally disinhibits the VNC rebound circuit and drives stronger activation of pIP10, resulting in a bias towards Pslow production. How TN1 and other song VNC neurons [5] coordinate the production of the two pulse types remains to be elucidated, but they must ultimately act via the ps1 motor neuron [6], shown to be required for males to switch from Pslow to Pfast when far from females [20].

Fourth, our study also provides a mechanistic explanation for a prior discovery of two hidden internal states in the male brain underlying song production, termed ‘close’ and ‘chasing’ [57]. Our work suggests that the P1a disinhibition arm of the pathway underlies the difference in these two states - in the ‘close’ state, in which sine song dominates and males are close to females, the P1a disinhibition circuit is engaged and sensory-driven pIP10 activity drives pulse-sine complex bouts. In the ‘chasing’ state, in which males are farther from females and moving faster, the P1a disinhibition circuit is not engaged and pIP10 activity drives primarily pulse-only simple bouts. This interpretation explains the observation that males continually toggle between ‘close’ and ‘chasing’ states throughout courtship, that ‘close’ state durations are longer than ‘chasing’ state durations, and why activation of pIP10 neurons in the presence of a female parodoxically both drove pulse song and pushed males into a state (’close’) that promoted sine song production [57].

Finally, our work also adds to the range of roles of the P1a neural cluster in modulating social behavior at different timescales [23, 25, 38, 39, 58]. While prior work emphasized the role of P1a in gating and sustaining male courtship behavior by controlling a minutes-long arousal state, here we identify an acute role for P1a in shaping behavior. We show that recent activation (timescales of milliseconds to seconds) of P1a neurons unlocks the potential for males to produce complex song (while separately, P1a neurons promote persistent singing). This may explain why males continually tap females throughout courtship (thereby directly activating P1a; [34, 59]): not only to maintain arousal, but to gate the production of long (complex) song bouts preferred by the female [54].

Our computational model of the song circuit reveals that a few key features (mutual inhibition, rebound excitability, and disinhibition) are sufficient, in combination with excitatory drive from fluctuating contextual cues, to recapitulate natural song dynamics (Fig. 5). These same features have been shown to contribute to motor pattern generation in both invertebrates and vertebrates [13, 42, 60, 61, 62, 63, 64, 65, 66, 67], although they are combined in new ways within the male song circuit. Such a minimalist circuit design both offers a simple control mechanism that allows the male to react to rapid changes in sensory context, and requires only few developmental changes to either derive this circuit from a unisex template [30] or alter the circuit to generate new song types in other species [21]. While we have not yet uncovered the identity of neurons mediating disinhibition downstream of P1a excitatory neurons or meditating the mutual inhibition within the song pre-motor circuit, new methods for large-scale imaging in *Drosophila* [68, 69, 70] and new connectomic resources for the *Drosophila* brain and VNC [41, 71, 72] will make it possible to discover these neurons and test their role in singing. Such work will ultimately be facilitated by our compact model of the song circuit.

## 8 Materials and Methods

### 8.1 Fly strains and rearing

See Table S2.

### 8.2 Behavioral apparatus

Behavioral experiments were performed in two custom made circular chambers (modified from [20]) within black acrylic enclosures. Ambient light was provided through an LED pad inside each enclosure (3.5” × 6” white, Metaphase Technologies). For each chamber, video was recorded at 60 fps (FLIR Blackfly S Mono 1.3 MP USB3 Vision ON Semi PYTHON 1300, BFS-U3-13Y3M-C, with TechSpec 25mm C Series VIS-NIR Fixed Focal Length Lens) using the Motif recording system and API (loopbio GmbH, Austria), and using infrared illumination of around 22 *μ*W/mm^2^ (Advanced Illumination High Performance Bright Field Ring Light, 6.0” O.D.,Wash Down, IR LEDs, iC2, flying leads) and an infrared bandpass filter to block the red light used for optogenetics (Thorlabs Premium Bandpass Filter, diameter 25 mm, CWL = 850 nm, FWHM = 10 nm). Sound was recorded at 10 kHz from 16 particle velocity microphones (Knowles NR-23158-000) tiling the floor of each chamber. Microphones were hand-painted with IR absorbing dye to limit reflection artifacts in recorded videos (Epolin Spectre 160). Temperature was monitored inside each chamber using an analog thermosensor (Adafruit TMP36).

### 8.3 Optogenetics

Flies were kept for 3-5 days on regular fly food or food supplemented with all-trans retinal (ATR) at 1 ml ATR solution (100 mM in 95% ethanol) per 100 ml of food. ATR-fed flies were reared in the dark. CsChrimson was activated at 1 *−* 205*μ*W/mm^2^, using 627nm LEDs (Luxeon Star).

### 8.4 Behavioral assays

For all behavioral experiments, virign males and virgin were used 3 to 5 days after eclosion. Experiments were started within 120 minutes of the incubator lights turning on. Males and females were single and group housed, respectively. Flies were gently loaded into the behavioral chamber before an experiment, using a custom-made aspirator. Females were placed first for paired experiments. Chamber lids were painted with Sigmacote (Sigma-Aldrich SL2) to prevent flies from walking on the ceiling, and kept under a fume hood to dry for at least 50 minutes prior to an experiment. Videos were manually scored for copulation. Data beyond copulation were excluded from analysis, unless statistical biases required exclusion of the entire recording.

#### 8.4.1 Free courtship

Free courtship recordings were performed for 30 minutes, as described in [18].

#### 8.4.2 Open loop neural activation

For open loop experiments, a fixed stimulus frequency of 1/8Hz was used. Stimulus irradiance could take four distinct values (0, 1, 25, 205*μ*W/mm2), spanning three orders of magnitude, and stimulus duty cycle could take five distinct values (1/64, 1/32, 1/16, 1/8, 2/8), and both irradiance and duty cycle were combined in a full factorial design, resulting in 16 distinct blocks (pooling blocks with zero irradiance) that were presented in pseudo-randomized order for 120 seconds each.

### 8.5 Offline song segmentation

For subsequent offline analysis, song was segmented as described previously [17, 20], using a modified sine detection parameter to account for different acoustics in the setup used here (`Params.pval` = 1*e* 7). For a given recording, the output of the song segmentation algorithm included information about the start and end of each bout and each sine train, as well as the center of each detected pulse, and a snippet of noise not including song. To reduce the risk of contaminating bout statistics with artificially split bouts due to low amplitude of sine song (the softer song mode), we excluded all bouts containing sine song with amplitude below a chosen signal-to-noise (SNR) threshold. Specifically, we estimated the noise amplitude using the noise segment that is automatically detected and returned by the song segmentation software (thus not containing song), by first reducing the 16-dimensional (for 16 microphones) noise segment to a one-dimensional vector by storing the noise value of the loudest microphone at each time point, and then defining noise amplitude as the 99th percentile of the absolute value of the one-dimensional noise vector. Sine amplitude was calculated similarly, such that the SNR for a given sine bout was the ratio of the sine amplitude and the noise amplitude. We excluded bouts containing sine song with an SNR below 1.3 from further analysis. Further, the song segmenter occasionally split individual sine trains, due to intermittent noise. Uncorrected, this could for example split a ‘psp’ bout into one ‘ps’ and one ‘sp’ bout very close in time. This allowed us to use a simple temporal threshold to merge such bouts if the inter-bout interval was below 0.5s. The segmentation software is freely available at https://github.com/murthylab/MurthyLab_FlySongSegmenter.

### 8.6 Tracking

Male and female pose (locations of head, thorax, left and right wing tip) was automatically estimated and tracked, and manually proofread for all videos using `SLEAP` ([27], `sleap.ai`).

### 8.7 Song behavior analysis

#### 8.7.1 Song probabilities

For experiments with open loop optogenetic neural activation, the probability for a male to sing pulse or sine song at any point in time during a trial of a given stimulus block was computed as the fraction of trials containing pulse or sine song. For analyses separating song probabilities into far and near contexts, the average male-female distance (mfDist) within a trial was thresholded to assign the trial to one of the two contexts. Song probabilities for each context were then calculated using only those trials assigned to that context.

#### 8.7.2 Song sequences

Song segmentation provided information about the start and end of each bout, and all pulse and sine events within a bout, allowing to assign each bout a label describing the sequence of contained pulse and sine trains (‘p’ for a bout containing only pulse song, ‘spspspsp’ for a bout starting with sine song followed by several alternations between pulse and sine). For statistics, we reduced the amount of different bout types by abbreviating all bouts with two or more song alternations as ‘ps…’ or ‘sp…’ and referred to these as ‘complex p’ or ‘complex s’. ‘Acute’ and ‘persistent’ bouts were defined as bouts starting during a stimulus or after stimulus offset, respectively. ‘Rebound’ song was defined as song that started after stimulus offset, in a bout that started during a stimulus (e.g., if the initial pulse train in a ‘ps’ bout starts during a stimulus, but the following sine train starts after stimulus offset, that is considered rebound sine).

#### 8.7.3 Tap detector model

The tap detector model was constructed using a convolutional neural network (CNN). The CNN consisted of two 2D convolutional layers followed by two fully connected layers. The two convolutional layers had 32 and 64 output channels, respectively, a kernel size of five, and a stride of 1. The outputs of each convolutional layer were passed through a rectified linear unit (ReLU) nonlinearity and a 2D max pooling layer with a kernel size of 2 and stride of 2. The first fully connected layer had 53824 input and 32 output features followed by a ReLU nonlinearity, and the second fully connected layer had 32 input and 2 output features corresponding to scores for a tap or non-tap. The model was trained using the AdamW algorithm for 100 epochs with a batch size of 16 and a learning rate of 0.0001. The model was constructed and trained using the PyTorch library [73].

To train the CNN, video frames (size 128×128) of courting flies centered on the male were manually labelled as a tap or non-tap event using a custom GUI. Ten videos were used for creating the tap dataset, with 12,606 manual annotations total. Of these annotated frames, 70% were used for training and 30% were held out for model validation. Receiver-operating characteristic analysis was performed on held out data to determine the relationship between model recall (true-positive rate) and fall-out (false-positive rate) as a function of tap-detection threshold.

#### 8.7.4 Tap-based model of P1a neural activity

We convolved the binary output of the tap detection network (tap=1 / no tap=0, using a threshold on tap probability of *p*(tap) ≥ 0.9) with the known calcium fluorescence of P1 neurons in response to a single tap of the female abdomen (tap-triggered average, [34]) to get an estimate of P1a neural activity in freely courting males on a moment-to-moment basis. We deconvolved the estimated calcium fluorescence signal with a kernel of the GCaMP6s calcium response (time constant 2.6s,[74]) to obtain an estimate of P1a rate, which we used for further analysis.

#### 8.7.5 Bout-triggered analysis of tap rate

For a given recording, the binary tap detector output at video resolution was first upsampled to audio resolution, using the camera trigger signal for synchronization. For each song bout with leading pulse song (simple ‘p’ or complex ‘ps…’), the number of detected taps occurring during the bout, *N*_during_, was counted, and this was divided by the duration of the bout, *B*, to produce tap rate *during* the bout, *R*_during_ = *N*_during_/*B*. As a control, the tap rate *before* the bout was computed as the number of taps occurring in an equally sized time window *B* immediately preceding the bout, *R*_before_ = *N*_before_/*B*. Tap rates were averaged (using the mean) per animal across simple and complex bouts and used for further analysis.

#### 8.7.6 Generalized linear model analysis

To estimate the relative predictive power of different sensory features on the choice of bout (here, complex vs. simple p), we used the generalized linear modeling framework with a sparse prior to penalize non-predictive history weights, as described before [18, 75]. Briefly, 10 sensory features (male and female forward velocity, mFV/fFV, lateral speed mLS/fLS, rotational speed, mRS/fRS, the angle of the male (female) thorax relative to the female (male) body axis, fmAngle / mfAngle, the distance between the male and female thorax, mfDist, and the instantaneous rate of P1a neurons estimated from detected taps, P1rate) were first smoothed using a moving average filter with a width of 20 video frames (0.33 seconds). Then, 21 uniformly distributed samples were extracted from the smoothed features within the five seconds of history leading up to the end of the first pulse train of each bout with leading pulse song (for simple pulse bouts, this corresponded to the end of the bout). Extracted features were z-scored per feature, to account for different feature dimensions and scales. Input to the GLM were the transformed features and a corresponding binary vector indicating whether a given feature history corresponded to a simple or complex pulse bout, and output were estimated filters for each feature (providing information on which dynamics in the feature, within the history window, were most predictive for bout type) and the relative deviance reduction (a measure of model performance). To estimate fit robustness, we repeated GLM fitting 51 times, each time using 70% of the input data (sampled randomly without replacement). For each feature, the mean across fits and the mean absolute deviation from the mean across fits were calculated and used for display.

### 8.8 Two-photon functional imaging

We imaged brain activity of Dsx+ cells in the VNC following pIP10 optogenetic activation using a custom-built two-photon laser scanning microscope [68, 76]. Virgin male flies (5-8 days old) were mounted and dissected as described previously [31], with minor differences. In brief, we positioned the fly ventral head and thorax side facing up to the underside of the dissection chamber, exposing both the ventral side of the central brain and the ventral side of the VNC. From the head, we removed the proboscis, surrounding cuticle, air sacks, tracheas, and additional fat or soft tissue. From the VNC, we removed thoracic tissue ventral to the VNC (e.g. legs and cuticle), exposing the 1st and 2nd segments of the VNC. Perfusion saline was continuously delivered to the meniscus between the objective and dissection chamber throughout the experiment. We imaged Dsx+ TN1 cells (1 hemisphere at a time), located in the ventral side of the 2nd segment of the VNC. We recorded 3-4 subvolumes of approximately 70 × 70 × 20 um3 at a speed of 1 Hz (0.3 × 0.3 × 2 um3 - 0.4 × 0.4 × 2 um3 voxel size) covering the full ventral-to-dorsal extent of the TN1 cluster ( 70 um). Volumetric data was processed using FlyCaIMan ([68]; https://github.com/murthylab/FlyCaImAn). In brief, volumetric time-series of GCaMP6s signal was motion-corrected in the XYZ axes using the NoRMCorre algorithm [77], and temporally resampled to correct for different slice timing across planes of the same volume, and to align timestamps of volumes relative to the start of the optogenetic stimulation (linear interpolation). Subvolumes consecutively recorded along the Z axis were stitched along the Z axis using NoRMCorre. Dsx+ TN1 somas were segmented by using the Constrained Nonnegative Matrix Factorization (CNMF) algorithm to obtain temporal traces and spatial footprints of each soma as implemented in CaImAn ([68, 78]; the initial number and XYZ location of all TN1 somas were manually pre-defined). For pIP10 activation, we used a stimulus of 2 seconds ON (at 13 uW/mm2 irradiance) and 2 seconds OFF repeated 4 times to maximize the magnitude of evoked GCaMP responses. Imaging started 10 seconds before stimulus onset, where baseline activity was measured, and lasted 10 seconds after stimulus offset.

### 8.9 Neural circuit model of song bout statistics

Network simulations were performed using the Brian2 package [79] with python3. Individual neurons were defined as variants of the Izhikevich model [43] with known spiking properties (such as rebound or tonic spiking) that matched experimental predictions. Briefly, the neuronal membrane potential *v* was modeled via

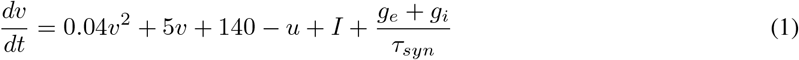

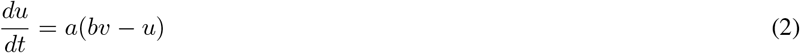

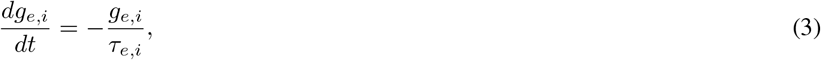

with the membrane recovery variable *u*, the timescale *a* and the sensitivity *b* to subthreshold fluctuations of the membrane potential of the recovery variable, and the input current *I*. *g*_*e*_ and *g*_*i*_ are excitatory and inhibitory conductances, and *τ*_*syn*_ is the synaptic time constant. Whenever the membrane potential reached 30mV, this was considered an action potential and the membrane variables were reset via

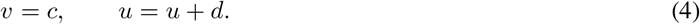

The full song circuit model comprised four Izhikevich neurons, termed ‘p’,‘s’ (for ‘pulse’ and ‘sine’), ‘pC2’, and ‘inh’. Parameters *a, b, c, d* were chosen to enable post-inhibitory rebound dynamics for the ‘p’ and ‘s’ node, and tonic spiking for the ‘pC2’ and ‘inh’ node (Table S3). Inhibitory connections were defined mutually between ‘p’ and ‘s’, from ‘inh’ to both ‘p’ and ‘s’, and from ‘pC2’ to ‘inh’. A single excitatory connection was defined from ‘pC2’ to ‘p’. For each spike in a presynaptic neuron, the synaptic conductance *g*_*e,i*_ was incremented by *w*_*e,i*_. *w*_*e,i*_ were free parameters that were fit during genetic algorithm optimization. The remaining free parameters were the amount of tonic input current into the ‘inh’ node (*I*_tonic_, regulating the amount of tonic inhibition onto the core ‘p’-‘s’ circuit), and a multiplicative factor *I*_*e*_ that controlled the gain of the sensory input current into ‘pC2’. The sensory input current into ‘pC2’ was the male-female distance (mfDist) during a given recording of wild-type courtship, subjected to nonlinear transformation via

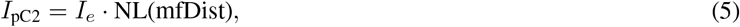

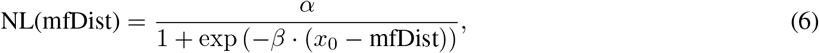

to facilitate strong/weak input current to pC2 at short/large distance. Numerical simulations of the network were performed using Euler integration, and spike times of each node were recorded for further analysis. Specifically, ‘song sequences’ of the model were defined based on the activity of the ‘p’ and ‘s’ node, such that a coherent spike train of one node that was at least 300ms separated from the next spike of the other node was considered a ‘simple’ bout, whereas alternating activity of the two nodes within 300ms was considered a ‘complex’ bout. All model parameters are specified in Table S4.

#### 8.9.1 Genetic algorithm optimization

The distribution of model bout types in response to a given naturalistic stimulus was directly comparable to the actual distribution of male song bouts corresponding to the sensory stimulus, which we exploited to fit the four free parameters of the model (a scalar gain factor for the input to the pC2 node, the strength of a constant input current to the inh node, and one global weight each for all excitatory and inhibitory connections) to the experimental data. Specifically, we used genetic algorithm optimization (the `geneticalgorithm` python package, https://pypi.org/project/geneticalgorithm/) to minimize the root mean squared difference (RMSD) between the experimental and simulated bout distribution (using six bout types, ‘p’,’ps’,’psp…’,’s’,’sp’,’sps...’, to provide more information to the algorithm than when using the four categories ultimately used for analysis; this led to slightly better model fits), as well as the absolute difference between the number of experimental and simulated bouts (∆*N*_bout_), via the objective function RMSD + 0.1 ∆*N*_bout_ (see Table S4 for optimization parameters and ranges). The relative scaling of the two objectives was chosen to prioritize reproducing the bout distribution over the number of bouts. All genetic algorithm parameters are specified in Table S4. 400-second pieces of song data, randomly chosen from all 20 wild-type recordings with at least 10% of song bouts produced far from the female (mfDist>4mm), were used as input to the genetic algorithm.

#### 8.9.2 Knockout simulations

To test the relevance of different computational features of the circuit model, we compared genetic algorithm fit performance for the full model (here using 200-second song snippets, randomly chosen from all wild-type recordings) to fit performance for versions of the model with individual computational features ‘knocked out’. Specifically, while in the full model both the p and s node were rebound excitable (by choosing the appropriate values for parameters *a, b, c, d* (see Table S3), rebound excitability was knocked out in the p (no rebound pulse), s (no rebound sine), or both nodes (no rebound) by adjusting parameters *a, b, c, d* (to turn these nodes from ‘rebound spiking’ into ‘tonic spiking’; see Table S3). Disinhibition was knocked out by removing the inhibitory synapses of the inh node onto the p and s nodes.

### 8.10 Irradiance measurements

Irradiance levels reported for optogenetic neural activation in freely behaving flies were measured (using a Thorlabs PM100D power meter) at the center of the experimental chamber, with the chamber lid in place. Two identical experimental setups were used for behavioral experiments, and irradiance levels were calibrated to have uniform voltage-to-irradiance conversion across setups.

Irradiance reported for optogenetic stimuli during two-photon calcium imaging was measured (also using a Thorlabs PM100D power meter) at approximately the level of the preparation (after the objective).

### 8.11 Statistics

Statistical analyses were performed either in Matlab 2019a or Python 3.7. The two-sided Wilcoxon rank-sum test (Mann-Whitney U test) for equal medians was used for statistical group comparisons unless noted otherwise. Errorbars indicate mean±mean absolute deviation (MAD) from the mean unless otherwise specified. Experimenters were not blinded to the conditions of the experiments during data collection and analysis. All attempts at replication were successful.

### 8.12 Data and code availability

Code will be available at github.com/murthylab and data will be available at `dataspace.princeton.edu`.

## 9 Acknowledgments

We thank Jesse Goldberg, Annegret Falkner, Jonathan Pillow, Ilana Witten, Jan Clemens, Sama Ahmed, and Bartul Mimica for comments on the manuscript. We thank Georgia Guan for technical assistance. We thank Minseung Choi for assistance with real-time song segmentation and development of closed-loop optogenetic protocols. Some figure schematics were created with BioRender.com.

## 10 Author Contributions

Data collection: Frederic A. Roemschied, Elise C. Ireland, Diego A. Pacheco, Xinping Li

Model and code development: Frederic A. Roemschied, Rich Pang, Max J. Aragon

Data analysis and generation of figures: Frederic A. Roemschied

Writing: Frederic A. Roemschied, Mala Murthy

Conceptualization: Frederic A. Roemschied, Mala Murthy

## 12 Materials and correspondence

Correspondence and material requests should be addressed to Mala Murthy.

