## Supplemental Information for "Flexible Circuit Mechanisms for Context-Dependent Song Sequencing"

### 1 Supplementary information

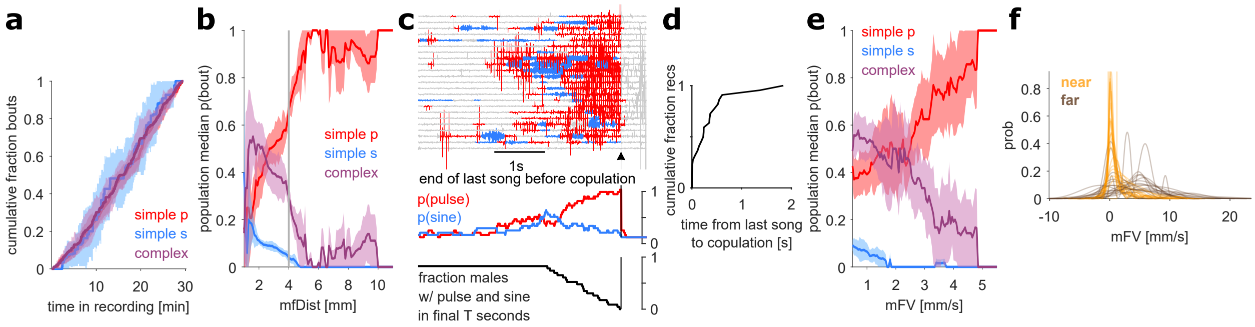


Figure S1: **Supplement to Fig. 1**
**a**, Cumulative fraction of simple pulse (red), simple sine (blue), or complex (purple) bouts over time in recording, for $n = 20$ wild-type male-female pairs.
**b**, Population-averaged probability of wild-type males singing simple pulse (red), simple sine (blue), or complex (purple) bouts at a given male-female distance (mfDist). Grey vertical line indicates the distance threshold of 4mm used to define far and near song bouts.
**c**, The majority (59%) of bouts immediately preceding copulation are complex ($n = 23$ wild-type pairs with copulation within a 20 minute recording). Song bouts are aligned to bout end. Time-resolved probability of pulse (red) and sine song (blue) (shown below song traces) rises prior to copulation. Black curve at the bottom shows the fraction of males that sing both pulse and sine song in the time prior to copulation. 80% of males sang both song modes within the final 1.5 seconds of song before copulation, suggesting complex bouts facilitate mating.
**d**, The majority (91%) of final bouts (the last song bout prior to copulation) occur within 0.6 seconds preceding copulation.
**e**, Population-averaged probability to sing simple pulse, simple sine, or complex bouts relative to male forward velocity (mFV). Color code as in (a).
**f**, Distribution of mFV near (yellow) and far (brown) from the female.
**a,b,e**, median±median absolute deviation from the median.
**a,b,e,f**, $n = 20$ recordings of male-female pairs.

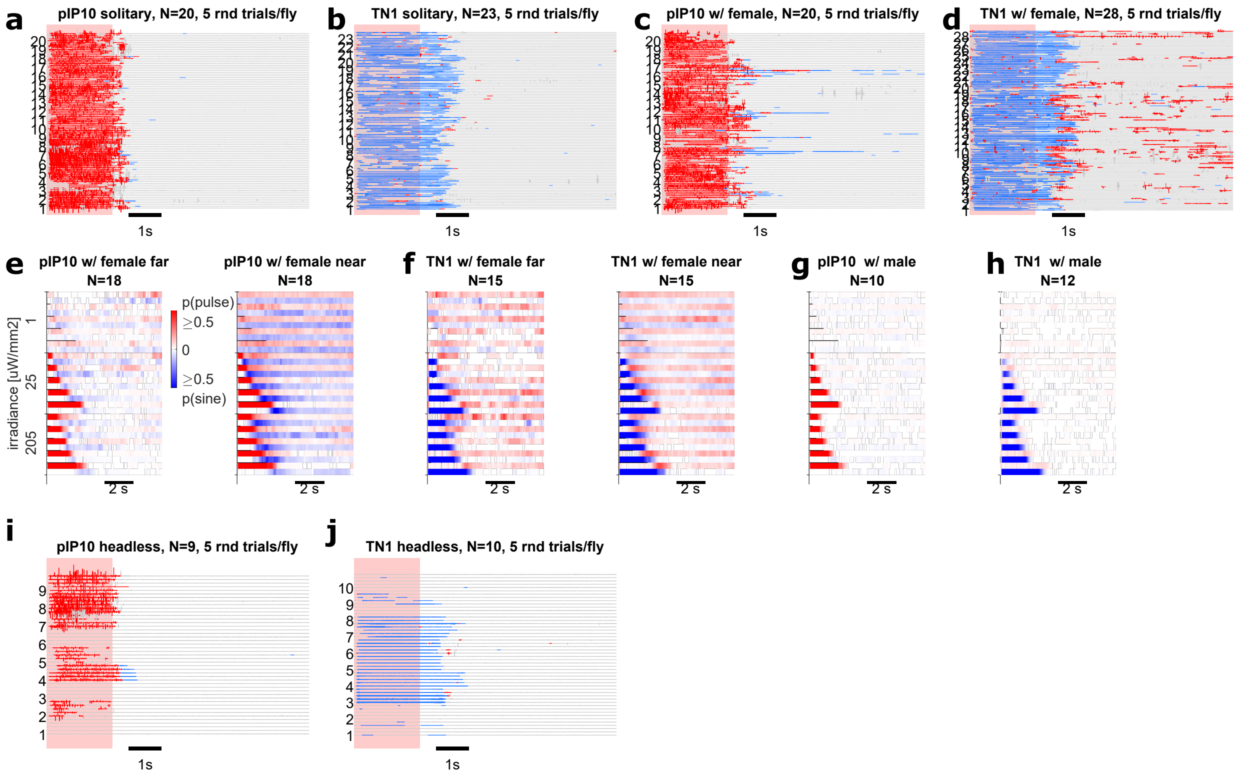


Figure S2: **Supplement to Fig. 2**
**a**, Example raw song responses drawn from $n = 20$ solitary pIP10>CsChrimson males with a single type of optogenetic stimulus (205uW/mm2 on for 2s per 8s trial). For every recording, five out of 15 trials were randomly chosen for display. Numbers on y-axis indicate recording. Color code: red - pulse song, blue - sine song, grey - silence, pink - optogenetic stimulus.
**b**, Example raw song responses drawn from $n = 23$ solitary TN1>CsChrimson males to the same stimulus type shown in a).
**c**, Example raw song responses frawn from $n = 20$ pIP10>CsChrimson males, paired with a wild-type female, to the same stimulus type shown in a).
**d**, Example raw song responses drawn from $n = 28$ TN1>CsChrimson males, paired with a wild-type female, to the same stimulus type shown in a).
**e**, Population-averaged song responses of $n = 18$ pIP10>CsChrimson males paired with a wild-type female as shown in Fig. 2d, but split into instances during which male and female were far or near (as quantified in Fig. 4e-f).
**f**, Population-averaged song responses of $n = 15$ TN1>CsChrimson males paired with a wild-type female as shown in Fig. 2i, but split into instances during which male and female were far or near (as quantified in Fig. 4e-f)..
**g**, Population-averaged song responses of $n = 10$ pIP10>CsChrimson males paired with a wild-type male.
**h**, Population-averaged song responses of $n = 12$ TN1>CsChrimson males paired with a wild-type male.
**i**, Example raw song responses drawn from $n = 9$ solitary headless pIP10>CsChrimson males to the same stimulus type shown in a).
**j**, Example raw song responses drawn from $n = 10$ solitary headless TN1>CsChrimson males to the same stimulus type shown in a).

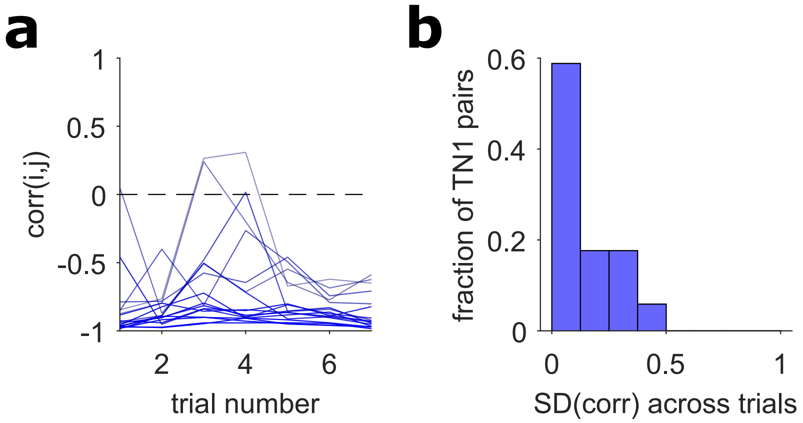


Figure S3: **Supplement to Fig. 3**
**a**, Anti-correlation between calcium responses of TN1 neuron pairs persists across trials. While Fig. 3c shows the correlation between trial-averaged calcium responses of TN1 neuron pairs in one fly, here we show the correlation between TN1 pairs for individual trials (7 trails, each trial consisting of four optogenetic stimulus presentations and a pause, as shown in Fig. 3b), only for pairs with trial-averaged anticorrelation coefficient below -0.8 ($n = 17$).
**b**, Standard deviation (SD) across trials of the correlation coefficients shown in a). The majority of anti-correlated TN1 pairs are consistent across trials.

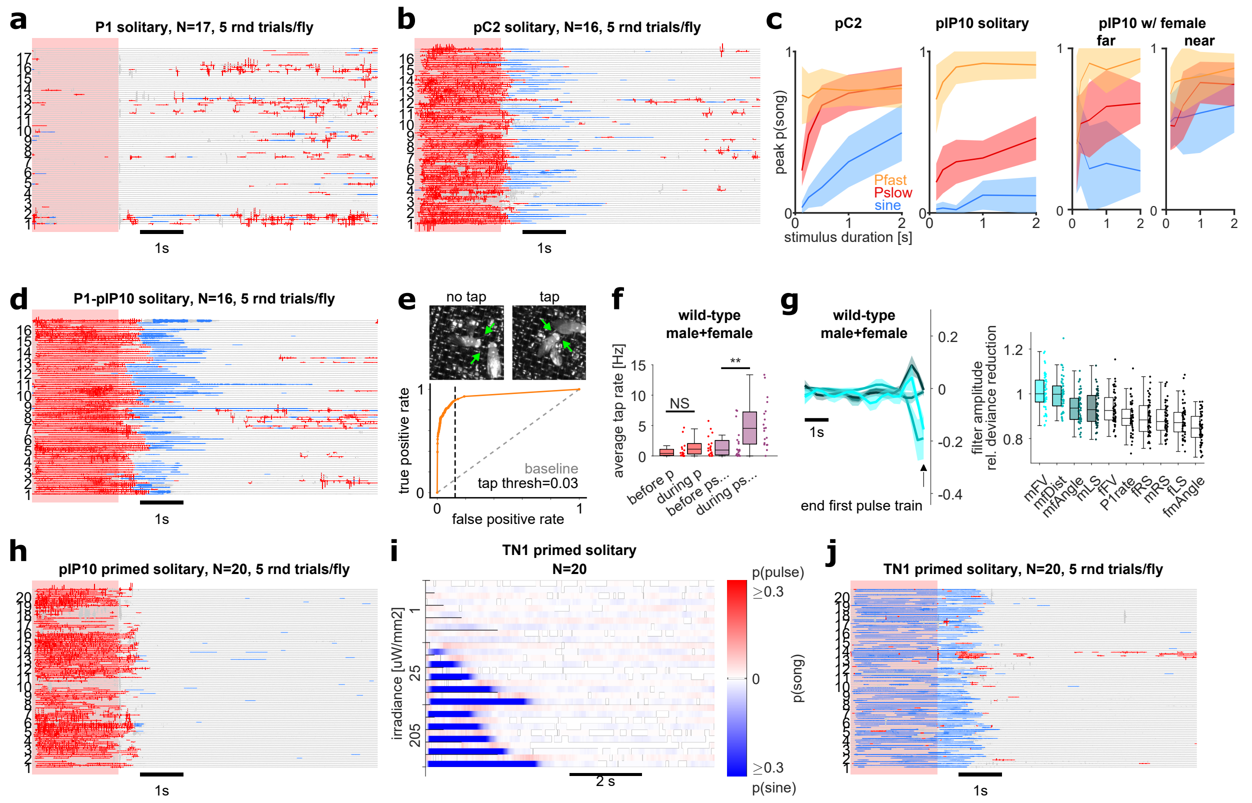

Figure S4: **Supplement to Fig. 4**

**a**, Example raw song responses drawn from $n = 17$ solitary P1a>CsChrimson males to a single type of optogenetic stimulis (205uW/mm2 on for 2s per 8s trial). For every recording, five out of 15 trials were randomly chosen for display. Numbers on y-axis indicate recording. Color code: red - pulse song, blue - sine song, grey - silence, pink - stimulus.
**b**, Example raw song responses drawn from $n = 16$ solitary pC2>CsChrimson males to the same stimulus type shown in a).
**c**, Peak probability of two types of pulse song termed Pfast and Pslow (orange and red; [1]) and sine song (blue) as a function of stimulus duration for intermediate-irradiance activation (25uW/mm2) of pC2 or pIP10 in solitary males, or pIP10 in males far or near from a wild-type female.
**d**, Example raw song responses drawn from $n = 16$ solitary P1a-pIP10>CsChrimson males to the same stimulus type shown in a).
**e**, Tap-detector model performance. (Top) Example of non-tap (left) and tap (right) events. Green arrows indicate the position of male foreleg tarsi. (Bottom) Receiver operator characteristic (ROC) curve for model after 100 epochs of training (orange points). Each point corresponds to a different tap probability threshold. Area under the ROC curve (AUC) is used as an evaluation metric – an ideal model would have an AUC of 1. Performance of a null model (gray diagonal line) is included for comparison.
**f**, Average tap rate before and during simple and complex pulse bouts, for $n = 20$ wild-type male-female pairs (analog to Fig. 4i). Wilcoxon rank-sum test for equal medians; **$P < 0.01$; NS, not significant.
**g**, A generalized linear model (GLM) to predict complex vs. simple pulse bout production based on the history of sensory features prior to the end of the first pulse train in each (ps... complex or p simple) bout in $n = 20$ recordings of wild-type male-female pairs (analog to Fig. 4k-l). Sensory features are ranked by their predictive power, and GLM filters are shown for the four most predictive features.
**h**, Example raw song responses drawn from $n = 20$ solitary pIP10>CsChrimson males to the same stimulus type shown in a). Males were primed (allowed to court a virgin wild-type female, to induce male courtship state) for five minutes prior to the start of the optogenetic stimulus protocol.
**i**, Population-averaged song responses of $n = 20$ primed solitary TN1>CsChrimson males.
**j**, Raw song responses of $n = 20$ primed solitary TN1>CsChrimson males to the same stimulus type shown in a).

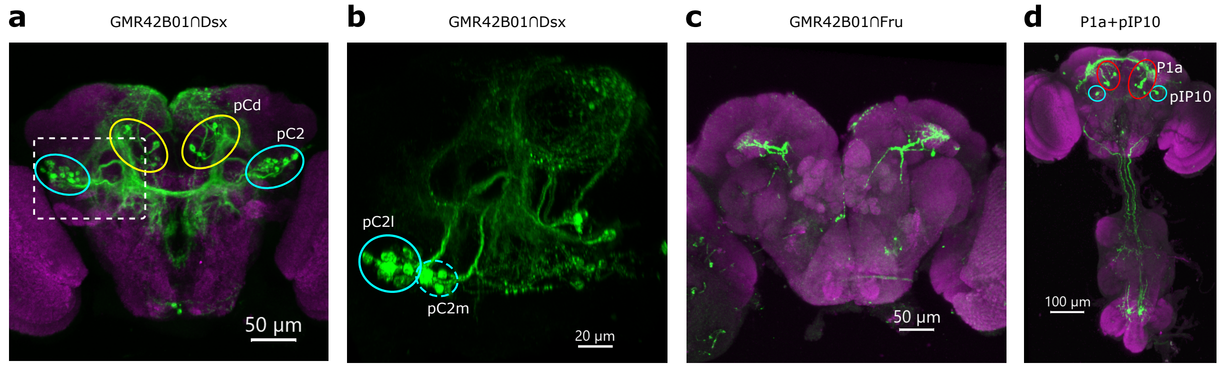


Figure S5: **Supplement to Fig. 4**
**a**, Male brain expressing CsChrimson.mVenus via GMR42B01 ∩ Dsx (green). Neuropil is labeled with nc82 (magenta). The intersection labels pC2 neurons (circled in blue), as well as 6 pCd-like neurons (circled in yellow). These pCd-like neurons do not express Fru (see panel c), and therefore constitute a different subset of neurons than the pCd neurons contributing to persistent male arousal downstream of P1a neurons in [2]. Broad-range optogenetic activation in males using the genetic driver for pCd neurons from [2] produces no song (data not shown).
**b**, Zoom of the boxed area in a), showing that the pC2 population labeled in the intersection consists of both pC2l neurons (solid circle) and pC2m neurons (dashed circle).
**c**, Male brain expressing CsChrimson.mVenus via GMR42B01 ∩ Fru (green). Neuropil is labeled with nc82 (magenta).
**d**, Male brain and VNC expressing CsChrimson.mVenus (green) via both P1a and pIP10 drivers. Neuropil is labeled with nc82 (magenta). Cell bodies of P1a are circled in red. Cell bodies of pIP10 are circled in cyan.
(see Table S2 for full genotypes)

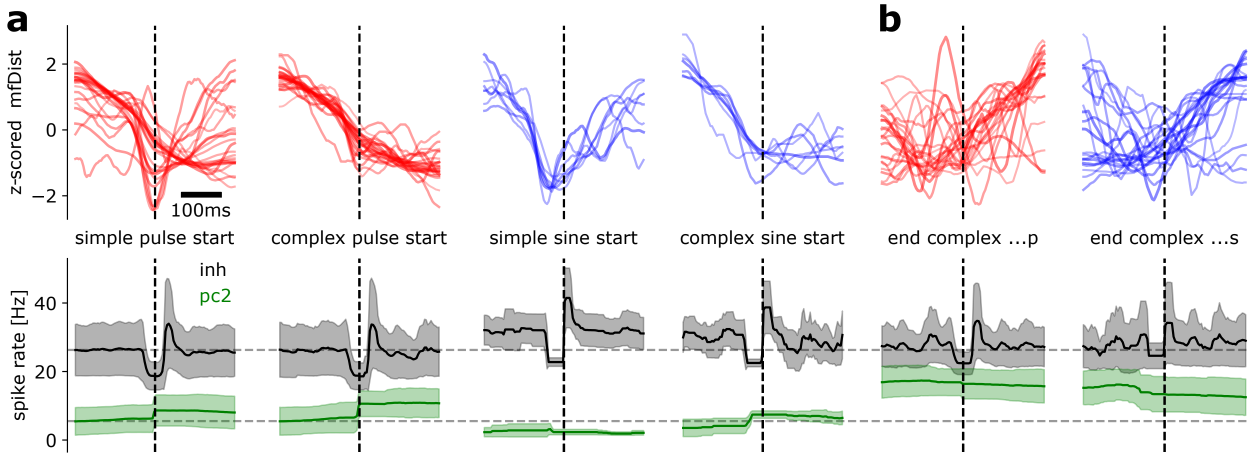


Figure S6: **Supplement to Fig. 5: Bout-triggered analysis in the song circuit model.**
**a**, (top) Z-scored male-female distance (mfDist) from wild-type courtship data (which served as input to the model) triggered around the time of simple (p,s) or complex (ps...,sp...) bout start in $n = 24$ simulations of the song circuit model. Each line is the z-scored mfDist averaged across bouts for one simulation (lines were smoothed for visualization, using a uniform filter of 44.4ms length). Every simulation uses 400 seconds of song randomly chosen from all wild-type recordings (such that the chosen song contained a minimum of 10% of bouts at mfDist $\geq$ 4mm, and the fit error / objective function value was below 0.1). For all bout types, mfDist decreases around the time of bout start. (Bottom) Instantaneous spike rate of the 'pC2' (green) and 'inh' (black) nodes of the circuit model around the time of bout onset. Distinctly timed release from inh-mediated inhibition in combination with distinct levels of pC2-mediated excitation drives different bout types.
**b**, Dynamics of z-scored mfDist (top) and instantaneous spike rate of the pC2 and inh nodes in the circuit model at the time of bout termination, for complex bouts ending in pulse (left) or sine mode (right). In both cases, bout termination is accompanied by increases in mfDist and a resulting reduction in pC2-mediated excitation of the pulse and sine node.

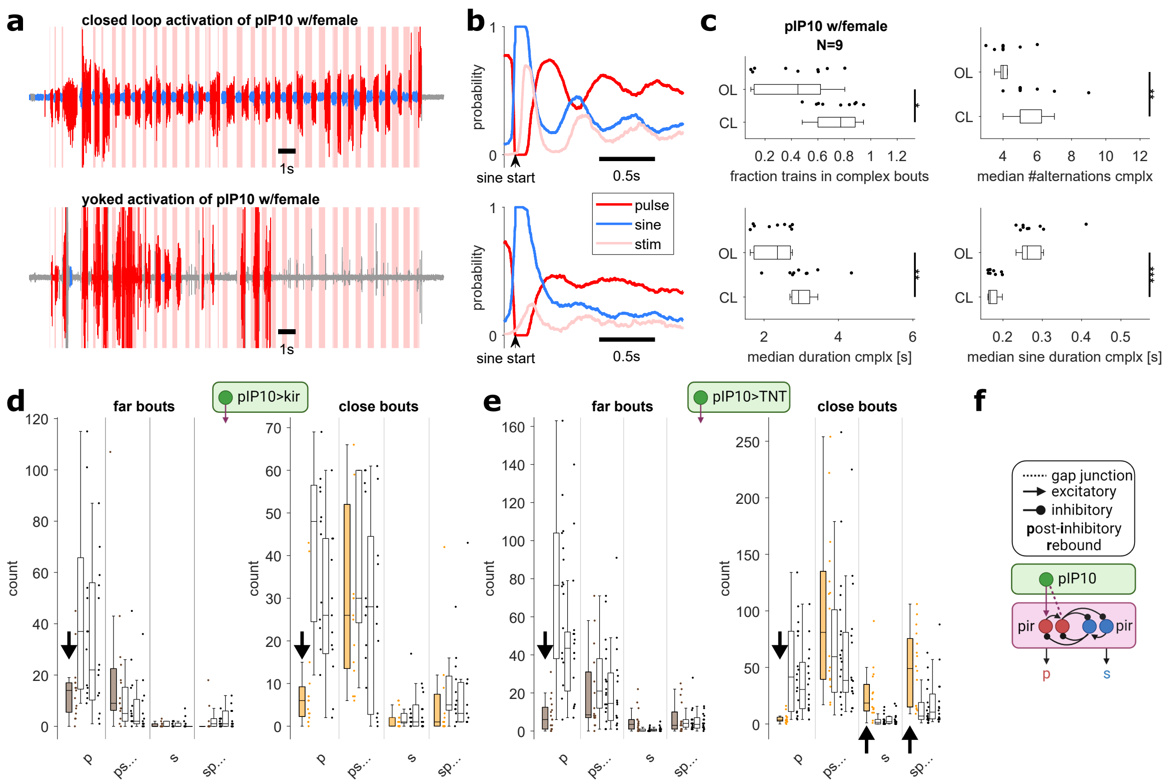


Figure S7: **Supplement to Fig. 5**

**a**, Triggering pIP10 activation on sine song in males courting a wild-type female strongly increases bout duration and complexity compared to controls with yoked activation (that is, identical stimulus statistics as in the closed loop condition but uncorrelated to the control male's song). Song shown from an example recording.

**b**, Song and stimulus probability around the onset of male sine song, for closed loop (top) and yoked (bottom) activation of pIP10 during the recording shown in a).

**c**, Population level comparison of four song features between closed-loop (CL) and yoked (OL) activation of pIP10: the fraction of trains belonging to complex bouts, the median number of sine-pulse or pulse-sine alternations in complex bouts, the median duration of complex bouts, and the median sine train duration within complex bouts. To show that all effects extend beyond generation of a single rebound sine, only 'psp...' bouts were considered for these analyses. Wilcoxon rank-sum test for equal medians; *P<0.05; **P < 0.01; ***P < 0.001.

**d**, Amount of pure and complex song bouts produced far from (brown) or near (orange) a female, in recordings of males with tonically hyperpolarized pIP10 neurons (via expression of inward-rectifying potassium channels in pIP10 neurons; VT040556>kir; filled box plots), compared to two genetic controls (blank box plots). See Table S2 for genotypes. The amount of pure pulse bouts was strongly reduced in males with hyperpolarized pIP10 both far from and near the female.

**e**, Amount of pure and complex song bouts produced far from (brown) or near (orange) a female, in recordings of males with blocked chemical synapses in pIP10 neurons (via expression of tetanus toxin light chain / TNT in pIP10 neurons; VT040556>TNT; filled box plots), compared to two genetic controls (blank box plots). The amount of pure pulse bouts was strongly reduced in males with blocked chemical synapses in pIP10 at both distances. Additionally, TNT males produced more bouts with leading sine song (s, sp...) compared to controls when near the female.

**f**, Circuit model to explain the findings in d-e: chemical synapses from pIP10 onto the VNC pulse node explain the reduction in pure pulse bouts with kir and TNT expression in pIP10. Gap junctions (electrical synapses) between pIP10 and the inhibitory interneuron node of the pulse pathway facilitate pure and complex sine bouts with blocked chemical synapses in pIP10, by transforming pIP10 activity through the electric synapses into inhibition onto sine driving neurons, leading to rebound sine bouts after termination of pIP10 activity.

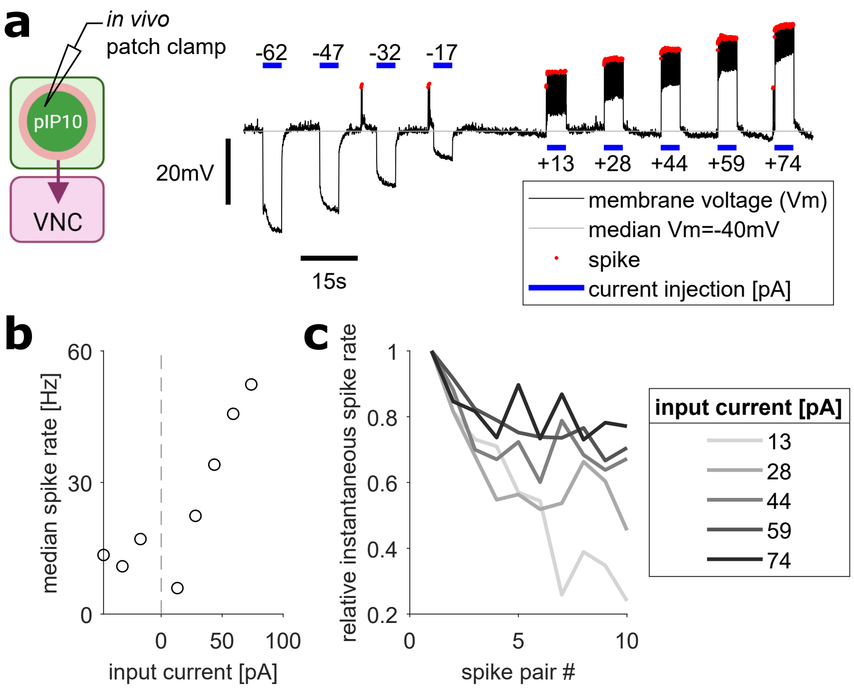


Figure S8: **Supplement to Fig. 5**
**a**, *In vivo* patch-clamp electrophysiology of descending neuron pIP10. Action potentials (spikes) were observed both during injection of positive current and following injection of negative current (post-inhibitory rebound spikes).
**b**, Median spike rate following negative or during positive current injection.
**c**, Instantaneous spike rate of the first ten spike pairs in each trial, normalized by the spike rate of the first pair. Normalized spike rate decreases for successive spikes, indicative of spike-frequency adaptation.

Supplementary Table S1: Key resources table

| Reagent type (species) or resource | Designation | Source or reference | Identifiers | Additional Information |
|---|---|---|---|---|
| Genetic reagent (*D. mel*) | NM91 | [3] | | A gift from Peter Andolfatto |
| Genetic reagent (*D. mel*) | 20xUAS-IVS-CsChrimson.mVenus (X) (attP18) | | BDSC: 55134 | A gift from Vivek Jayaraman and Gerry Rubin |
| Genetic reagent (*D. mel*) | VT040556-p65ADZp(attp40) | [4] | | A gift from David Stern |
| Genetic reagent (*D. mel*) | VT040347-ZpGDBD(attp2) | [4] | | A gift from David Stern |
| Genetic reagent (*D. mel*) | GMR15A01-AD(attp40) | [5] | | A gift from David Anderson |
| Genetic reagent (*D. mel*) | GMR71G01-DBD(attp2) | [5] | | A gift from David Anderson |
| Genetic reagent (*D. mel*) | GMR42B01-Gal4(attP2) | [6] | | A gift from Bruce Baker |
| Genetic reagent (*D. mel*) | dsx-LexA | [6] | | A gift from Bruce Baker |
| Genetic reagent (*D. mel*) | 8xLexAop2-flp | [6] | BDSC 55819-20 | Obtained from BDSC |
| Genetic reagent (*D. mel*) | R13H01.LexA::p65 (attP40) | [7] | | A gift from Troy Shirangi |
| Genetic reagent (*D. mel*) | dsxGal4(Δ2) | [8] | | A gift from Troy Shirangi |
| Genetic reagent (*D. mel*) | +/+; fruFLP/VT040556-Gal4 | [1] | | A gift from Barry Dickson |
| Genetic reagent (*D. mel*) | GMR53G02-AD (attP40)/UAS-Kir2.1 | [1] | | GMR53G02-AD(attP40) obtained from BDSC and contributed by Gerry Rubin |
| Genetic reagent (*D. mel*) | GMR53G02-AD (attP40)/UAS-TNT | [1] | | |
| Genetic reagent (*D. mel*) | UAS>stop>Kir2.1/+; fruFLP/VT040556-Gal4 | [1] | | UAS>stop>Kir2.1 was provided by Troy Shirangi |
| Genetic reagent (*D. mel*) | UAS>stop>TNT/+; fruFLP/VT40556 | [1] | | |
| Genetic reagent (*D. mel*) | norpA[36]-20xUAS-CsChrimson.mVenus(attP18);VT040556-AD/GMR15A01-AD;VT043047-DBD/GMR71G01-DBD | | | |
| Genetic reagent (*D. mel*) | ;;13xLexAop-IVS-GCAMP6s (attp1) | | BDSC 44273 | |
| Genetic reagent (*D. mel*) | ;LexAop-tdTomato.Myr, su(Hw) (attp5), BRP>STOP>V5-2A-LexA-VP16 [VK00018]/Cyo; TM2/TM6B, tb | | BDSC 56142 | |
| Chemical compound, drug | All-trans retinal | Sigma-Aldrich | #R2500 | |

Supplementary Table S2: List of strains used in manuscript

| Figure panel | Genotype | Additional Information |
|---|---|---|
| Fig. 1; Fig. S1a-c; Fig. S4f,g | NM91 | D. melanogaster NM91 males (provided by Peter Andolfatto) courting NM91 wild type females. |
| Fig. 2b-g; Fig. 4e,f,h-l,n-p; Fig. S2a,c,e,g,i; Fig. S4c,h; Fig. S7a-c | UAS-CsChrimson/+; VT040556-AD (attp40)/+; VT040347-DBD (attp2)/+ | Express CsChrimson in pIP10 neurons; VT040556-AD(attp40) and VT040347-DBD(attp2) [4] provided by David Stern |
| Fig. S8a-c | norpA[36]-20xUAS-CsChrimson.mVenus/+; VT040556-AD (attp40)/+; VT040347-DBD (attp2)/+ | Express CsChrimson in pIP10 neurons of norpA-blind flies; VT040556-AD(attp40) and VT040347-DBD(attp2) [4] provided by David Stern |
| Fig. S7d | UAS>stop>Kir2.1/+; fruFLP/VT040556-Gal4 | Express kir2.1 in pIP10 neurons [1] |
| Fig. S7d | GMR53G02-AD(attP40)/UAS-Kir2.1 | control kir2.1 |
| Fig. S7d,e | +/+; fruFLP/VT040556-Gal4 | pIP10 control |
| Fig. S7e | UAS>stop>TNT/+; fruFLP/VT040556-Gal4 | Express TNT in pIP10 neurons |
| Fig. S7e | GMR53G02-AD(attP40)/UAS-TNT | control TNT |
| Fig. 2h-l ; Fig. S2b,d,f,h,j; Fig. S4i,j | +/+; GMR13H01-LexA/20xUAS>stop>CsChrimson.mVenus(attP40); Dsx-Gal4/8xLexAop-FLP | Express CsChrimson in TN1 neurons; GMR13H01.LexA::p65 (attP40) and dsx-Gal4(Δ2) [7] provided by Troy Shirangi |
| Fig. 4b,e,f; Fig. S4b,c | UAS > stop > CsChrimson.mVenus(attP14)/8XLexAop2-flp; dsx-LexA, 8xLexAop2-flp/GMR42B01-Gal4 | Express CsChrimson in Dsx+ pC2 neurons [6] |
| Fig. 4d-f; Fig. S4d | norpA[36]-20xUAS-CsChrimson.mVenus(attP18);VT040556-AD/GMR15A01-AD;VT043047-DBD/GMR71G01-DBD | Express CsChrimson in pIP10 and P1a neurons |
| Fig. 4a,e,f; Fig. S4a | UAS-CsChrimson(attP18)/+; GMR15A01-AD (attp40)/+; GMR71G01-DBD (attp2)/+ (GMR15A01-AD (attp40) | Express CsChrimson in P1a neurons; GMR71G01-DBD (attp2) [5] kindly provided by David Anderson. |
| Fig. 3 | w[*] NorpA[36], 20xUAS-CsChrimson-mVenus (attp18); VT040556-AD(attp40)/ LexAop-tdTomato.Myr, su(Hw) (attp5), BRP>STOP>V5-2A-LexA-VP16 [VK00018]; VT400347-DBD (attp2)/Dsx-FLP, 13xLexAop-IVS-GCAMP6s (attp1) | Express mVenus-tagged CsChrimson in pIP10 neurons and GCaMP6s in Dsx+ neurons of norpA-blind males; |
| Fig. S8 | norpA[36]-20xUAS-CsChrimson.mVenus(attP18);VT040556-AD/+;VT043047-DBD/+ | Express mVenus-tagged CsChrimson in pIP10 neurons of norpA-blind males |

Supplementary Table S3: Izhikevich neuron parameters

| Neuron | a | b | c | d | Additional Information |
|---|---|---|---|---|---|
| pC2,inh | 0.02/ms | 0.2/ms | -65mV | 6V/s | 'tonic spiking' neuron |
| p,s | 0.03/ms | 0.25/ms | -60mV | 4V/s | 'rebound spiking' neuron |

Supplementary Table S4: Circuit model and genetic algorithm parameters

| Parameter | Value/Range | Additional Information |
|---|---|---|
| $\tau_i$ | 10ms | inhibitory time const. |
| $\tau_e$ | 2ms | excitatory time const. |
| $\tau_m$ | 5ms | membrane time const. |
| vth | 30mV | spiking threshold |
| v0 | -87mV | initial membrane potential |
| $\alpha$ | 1 | input nonlinearity |
| $\beta$ | 0.5 | input nonlinearity |
| $x_0$ | 2 | input nonlinearity |
| dt | 0.1ms | Euler integration time step |
| $I_e$ | $[2, 25]$ | Range used for optimization |
| $w_i$ | $[-150, -5]$ | Range used for optimization |
| $w_e$ | $[5, 150]$ | Range used for optimization |
| $I_{\text{tonic}}$ | $[3, 15]$ | Range used for optimization |
| number of iterations | 25 | |
| population size | 30 | |
| mutation probability | 0.1 | |
| elite ratio | 0.01 | |
| crossover probability | 0.5 | |
| crossover type | uniform | |

### 1.1 Supplemental Methods

#### 1.1.1 Fast online sine segmentation

To facilitate closed-loop optogenetic activation triggered on male sine song production, we used a custom-made fast online sine song segmenter that was based on two convolutional neural networks (CNNs). Briefly, at every time point during a recording, 384 samples (38.4ms) of sound history (384x16 samples for 16 microphone channels) served as input to the online segmenter. First, to reduce data dimensionality, the channel with the maximum mean power was selected for further processing and normalized by dividing by its 2-norm. We then applied a 512-point discrete Fourier transformation (DFT) to the normalized 1-dimensional signal and kept the 51 DFT amplitudes for frequencies below 1000 Hz for further processing. The normalized 384-sample time-domain waveform and the 51-sample frequency-domain amplitudes were fed into a classifier model that combined two CNNs. The output of the model was a sigmoidal activation value associated with the probability that the input data contained sine song. We used a threshold of $0.995$ on this activation value to create a binary output value that controlled the optogenetic stimulus LEDs during closed-loop neural activation (turning the stimulus on for $p(\text{sine}) \geq 0.995$ and off for $p(\text{sine}) < 0.995$). The CNNs were implemented using Keras [9] with Theano backend [10]. The model was trained on output of the offline song segmenter for $n = 19$ recordings of wild-type (NM91) male-female pairs not used otherwise in the present study. During these recordings, optogenetic stimulus LEDs were randomly turned on and off to generate a realistic noise background (as during closed-loop neural activation). Segmented song split up into 384x16-sample chunks (window size x number of microphones) and each chunk was labeled as ‘sine’ or ‘no sine’ to provide ground truth data for training. The recordings were divided into 11, 4, and 4 recordings for training, validation, and evaluation, respectively. During training, class weights were applied such that the less-represented class would have, on average, equal loss as the over-represented class in a completely naive, untrained classifier.

#### 1.1.2 Closed-loop neural activation

For closed-loop neural activation, stimulus LEDs were turned on at $51.5\mu\mathrm{W/mm2}$), triggered on the detection of sine song, using fast online sine segmentation (see above). For yoked control experiments, the stimulus of a closed-loop experiment was routed to the second chamber, such that stimulus statistics were identical between experiments but the correlation to male behavior was lost in the yoked control. In each experiment, the two chambers were randomly chosen for closed-loop activation or yoked control.

#### 1.1.3 Patch clamp recordings

Newly eclosed CsChrimson-expressing male flies were collected and raised on food that contained all-trans retinal for 24h before experiments. Flies were anesthetized on ice and mounted on a fly holder using ultraviolet glue. The cell bodies of pIP10 were made accessible by removing part of the cuticle, some trachea, and the perineural sheath covering the posterior brain. During the *in vivo* patch-clamp experiment, the fly dorsal head was continuously perfused with extracellular saline (103mM NaCl, 3mM KCl, 5mM N-Tris (hydroxymethyl)methyl-2-aminoethane-sulfonic acid, 8mM trehalose, 10mM glucose, 26mM NaHCO3, 1mM NaH2PO4, 1.5mM CaCl2 and 4mM MgCl2, 275–280mOsm, pH 7.3) bubbled with 95% O2 and 5% CO2. pIP10 neurons were identified as fluorescent from the mVenus and were recorded on either side of the brain, using glass electrodes with tip-resistances between 5–7 MΩ, filled with intracellular saline (140mM potassium aspartate, 10mM HEPES, 5mM EGTA, 4mM MgATP, 0.5mM Na3GTP, 1mM KCl, adjusted to 260–270mOsm, pH 7.3). Negative and positive current pulses of five seconds (with ten seconds between pulses) were injected into pIP10. Recordings were obtained using a MultiClamp 700B amplifier and then digitized with a NiDAQ PCI-6251 (National Instruments) at 10 kHz. Corrections of liquid junction potential correction were not performed.

#### 1.1.4 Immunohistochemistry

Fly dissections were performed on 3-6-day-old adult males in chilled phosphate-buffered saline (PBS). The dissected brains and VNSs were fixed in 2% paraformaldehyde in PBS with 0.1% Triton X-100 (PBT) for 55 minutes at room temperature and then were washed ten times for 5 min each with 0.5% PBT. Following fixation, brains and VNCs were moved to a blocking solution of 5% normal goat serum in 0.1% PBT for 2h at room temperature and then were transferred to a primary antibody solution for incubation for 3d at 4°C. The primary antibody solution contains mouse anti-Bruchpilot (nc82, 1:30, Developmental Studies Hybridoma Bank) and chicken anti-GFP (1:1,000, 1:1000) diluted in the blocking solution. After primary incubation, brains and VNCs were washed ten times for 5 min each with 0.5% PBT before they were moved to a secondary antibody solution for incubation for 4 hr at room temperature and then 24 hr at 4°C. The secondary antibody solution contains Alexa 488-conjugated goat anti-chicken (1:250, Invitrogen) and Alexa 568-conjugated goat anti-mouse (1:250, Invitrogen) diluted in the blocking solution. After ten final washes for 5 min each with 0.5% PBT, brains and VNCs were mounted on slides in Vectashield (Vector Laboratories) for confocal imaging.

Brains and VNCs were imaged using a Leica confocal microscope (TCS SP8) with either a Leica HC PL APO 20x/0.75 CS2 objective or a Leica HC PL APO 63x/1.40 Oil CS2 objective. Maximum intensity projections were generated from stacks of optical sections (taken at 0.69-2um spacing) using Imaris 9.8 (Oxford Instruments).

#### 1.1.5 Pulse type classification

As part of offline song segmentation, detected pulses were classified into two pulse types (Pfast and Pslow) using the methodology described in [1]. Briefly, the pulse classifier takes as input the pulse waveforms returned by the offline song segmenter and assigns pulse type based on the similarity of the pulse waveform to templates for each pulse type.
